# Alternative translation initiation produces synaptic organizer proteoforms with distinct localization and functions

**DOI:** 10.1101/2024.02.16.580719

**Authors:** Paul Jongseo Lee, Alexa R. Soares, Yu Sun, Caroline Fai, Marina R. Picciotto, Junjie U. Guo

## Abstract

While previous studies suggest that many mRNAs contain more than one translation initiation site (TIS), the biological significance of most alternative TISs and their corresponding protein isoforms (proteoforms) remains undetermined. Here we show that alternative translation initiation at a CUG and an AUG TIS in neuronal pentraxin receptor (NPR) mRNA produces two proteoforms, and their relative abundance is regulated by both neuronal activity as well as an adjacent RNA secondary structure. Downstream AUG initiation transforms the N-terminal transmembrane domain into a signal peptide, thereby converting NPR to a secreted factor sufficient to promote synaptic clustering of AMPA-type glutamate receptors. Changing the relative proteoform ratio, but not the overall NPR abundance reduces AMPA receptor in parvalbumin (PV)-positive interneurons and induces changes in learning behaviors in mice. In addition to NPR, N-terminal extensions of C1q-like synaptic organizers, mediated by upstream AUU start codons, anchor these otherwise secreted factors to the membrane. Thus, our results uncovered the plasticity of N-terminal signal sequences regulated by alternative TIS usage as a widespread mechanism to diversify protein localization and functions.

## INTRODUCTION

Eukaryotic mRNAs are predominantly monocistronic in nature. After scanning from the 5ʹ end of mRNAs, ribosomes initiate translation often at the 5ʹ-most AUG start codon, translocate by one codon at a time through the open reading frame (ORF), and terminate at the first in-frame stop codon, producing one polypeptide chain (Kapp and Lorsch, 2004). Deviations from this conventional model such as internal ribosomal entry sites (Shatsky et al., 2018), ribosomal frameshifting during elongation (Atkins et al., 2016), and stop-codon read-through (Loughran et al., 2014) had been considered either rare exceptions or restricted to viral mRNAs. In contrast, high-throughput translatomics studies such as those using ribosome profiling (Ingolia et al., 2011; Lee et al., 2012) have identified numerous previously unannotated ORFs, many of which can be validated by proteomics and when disrupted can cause detectable cellular phenotypes (Chen et al., 2020). While these studies suggest a large contribution of alternative ORFs to proteome diversity, the functions of the vast majority of the alternative translation products are not known. A large category of alternative ORFs initiate at alternative TISs that are in-frame with the canonical TISs and are expected to produce either N-terminal extended or truncated proteoforms (Ivanov et al., 2011; Van Damme et al., 2014). To date, only a few N-terminal proteoforms have been functionally characterized (Bogaert et al., 2020; Calligaris et al., 1995; Kazak et al., 2013; Tsang and Cheeseman, 2023; Williams et al., 2014), with a generalizable mechanism by which alternative TIS usage regulates protein functions still lacking.

Initially identified as binding proteins for the snake venom toxin taipoxin, neuronal pentraxins are a family of synaptic organizers that have been shown to recruit AMPA receptors to excitatory synapses at least in part by direct protein-protein interactions between their pentraxin domain and the N-terminal domains of AMPA receptor subunits. Neuronal pentraxin 1 (NP1) and 2 (NP2, also known as neuronal activity-regulated pentraxin or NARP), encoded by *NPTX1* and *NPTX2* genes, respectively, are secreted factors with N-terminal cleavable signal peptides. A third member in this family, neuronal pentraxin receptor (NPR), encoded by the *NPTXR* gene, is a type II membrane protein with a single N-terminal transmembrane domain that also functions as a signal anchor that targets the ribosome/mRNA complex to the endoplasmic reticulum (ER) surface. Intriguingly, an early study of NPR has found that the initial methionine residue corresponds to a CUG TIS in its mRNA sequence (Dodds et al., 1997). This CUG codon in NPR mRNA is highly conserved among vertebrates (**Figure 1A**), hinting at potential unique functions that cannot be readily fulfilled by an AUG TIS.

**Figure 1.**
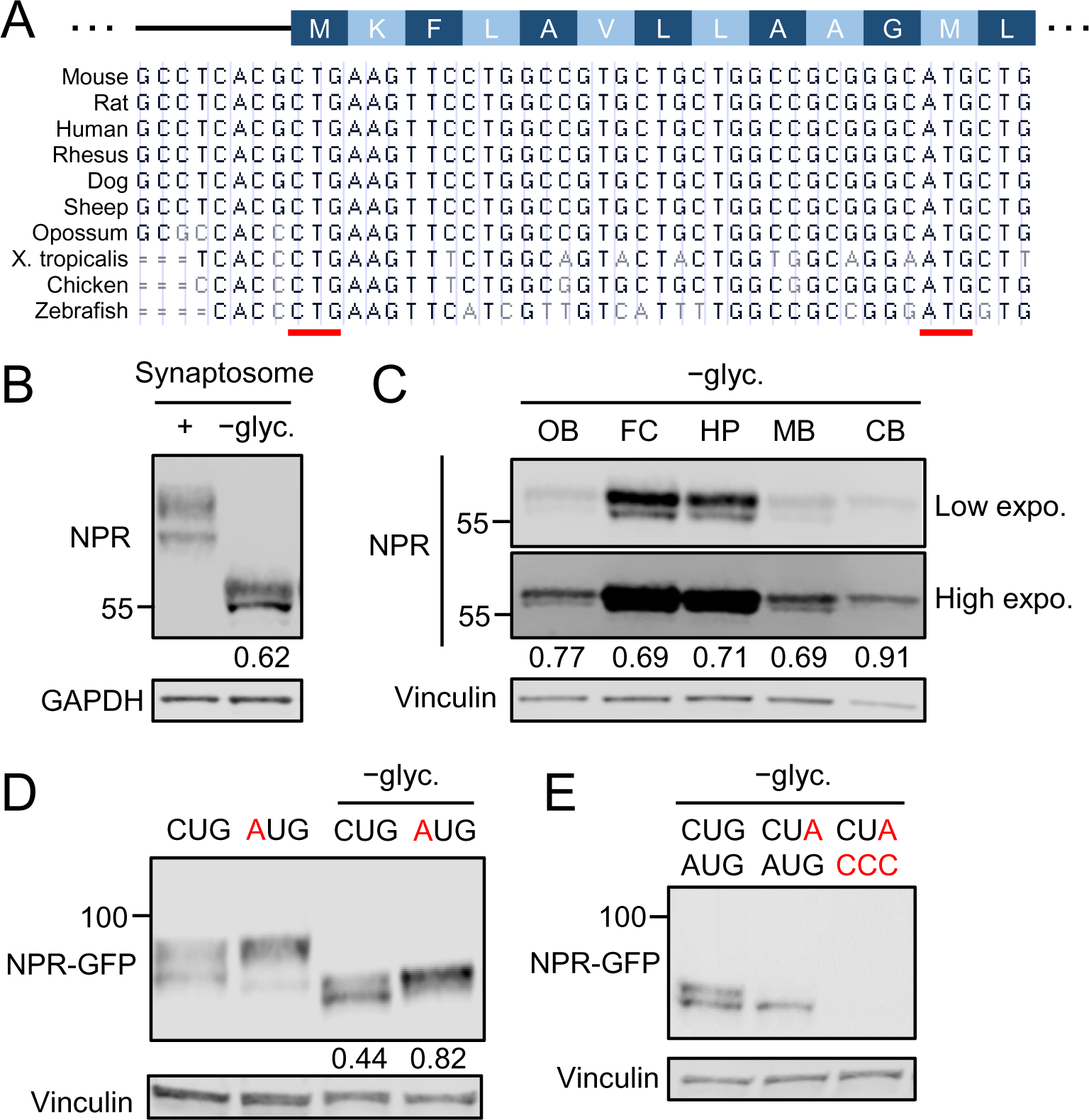
Alternative translation of NPR mRNA produces two N-terminal proteoforms. (**A**) Peptide sequence (top) and genome alignment (bottom) of the NPR N-terminal region across vertebrates. The annotated CUG TIS and downstream AUG TIS are underlined (red). (**B**) Endogenous NPR expression in forebrain mouse synaptosomes with or without glycans (−glyc). Long NPR fractions are shown below the immunoblots. (**C**) Endogenous NPR expression in different brain regions (OB: olfactory bulb, FC: frontal cortex, HP: hippocampus, MB: midbrain, CB: cerebellum). (**D**) Ectopic expression of wild-type NPR (CUG) and mutant (AUG) NPR cDNA in HEK293T cells. (**E**) Ectopic expression of NPR cDNAs with TIS mutations. All samples were deglycosylated in subsequent Western blot analyses unless otherwise indicated.

Here, by investigating the functional role of this noncanonical TIS in NPR mRNA, we uncovered a widespread mechanism by which alternative TIS usage switches the function of N-terminal signal sequences and the localization of proteoforms. Leaky scanning at the CUG TIS followed by the alternative translation initiation at a downstream AUG TIS remodels the N-terminal signal anchor to a cleavable signal peptide, thereby converting transmembrane NPR to a secreted, short NPR proteoform with distinct localization and function. A CTG-to-ATG knock-in mouse model with an altered NPR proteoform ratio showed reduced AMPA receptor abundance in PV^+^ interneurons and changes in learning behaviors. We identified dozens of additional dual proteoforms encoded by mRNAs from a wide variety of biological pathways including all four members of the C1q-like (C1QL) family of synaptic organizers, suggesting widespread impact of alternative TIS usage on protein localization and functions in cells.

## RESULTS

### Alternative translation of NPR mRNA produces two N-terminal proteoforms

As NPR has been shown to primarily regulate excitatory synapses (Cho et al., 2008; Lee et al., 2017; Pelkey et al., 2015), we analyzed its endogenous expression in mouse forebrain synaptosomes. As expected for most membrane proteins and consistent with previous studies (Dodds *et al*., 1997), NPR is glycosylated at multiple asparagine residues (**Figure 1B**). After the removal of glycans by a mix of *N*- and *O*-glycosidases, two distinct species with slightly different molecular weights were still observed (**Figure 1B**). The two species were readily detectable in most brain regions, with the exception of the cerebellum, in which NPR abundance was low and the shorter species was less apparent (**Figure 1C**). To test whether the distinct NPR species may be related to the CUG start codon, we ectopically expressed either wild-type (WT) NPR cDNA along with its native 5ʹ untranslated region (UTR), or a mutant with the CUG TIS replaced by an AUG. Similar to endogenous NPR, WT NPR cDNA also produced two proteoforms in both HEK293T cells (**Figure 1D**) and mouse primary cortical neurons (**Figure S1**), although the ratio between the two proteoforms appeared to be skewed towards the long form in neurons (73% versus 44% in HEK293T cells). NPR mRNA with an AUG TIS produced predominantly the long form (**Figure 1D, Figure S1**), indicating that the short NPR proteoform was not due to post-translational modification nor partial degradation from the long proteoform, and that its expression is specific to WT NPR mRNA with a CUG TIS.

The TIS-dependent proteoform heterogeneity suggested that they may be due to alternative translation initiation. A mutant NPR mRNA with the first 11 codons deleted (ΔN) produced an N-terminal truncated product with a similar size to that of the short NPR (**Figure S1**), consistent with the possibility of the downstream AUG being an alternative TIS. Moreover, substituting CUG with CUA eliminated the long but not short NPR proteoform (**Figure 1E**), while additionally mutating the downstream AUG further eliminated short NPR (**Figure 1E**), confirming that it was indeed the product of this alternative TIS. Therefore, both endogenous and ectopically expressed NPR mRNAs undergo alternative translation initiation at either the previously annotated CUG TIS or the downstream AUG, producing two N-terminal proteoforms.

### A stable RNA secondary structure promotes CUG initiation in mammals

Based on genome alignments, both the CUG and downstream AUG TISs are conserved across vertebrates (**Figure 1A**). To test whether the alternative production of dual proteoforms is also conserved, we *in vitro* transcribed four reporter mRNAs, each of which contains the first 135 nucleotides of NPR coding sequence from either human, mouse, frog, or zebrafish, followed by a GFP sequence, and translated each mRNA in rabbit reticulocyte lysates at the corresponding body temperature (see Methods). Both proteoforms were produced from each of the four NPR orthologs (**Figure 2A**), suggesting that alternative translation initiation is indeed evolutionarily conserved. Interestingly, however, the CUG initiation efficiency was much higher in human and mouse than that in frog and zebrafish NPR mRNAs (**Figure 2A**). Because all four reporter mRNAs were translated in the same *in vitro* system, the difference in CUG initiation efficiency must be due to cis-regulatory elements that control proteoform ratios.

**Figure 2.**
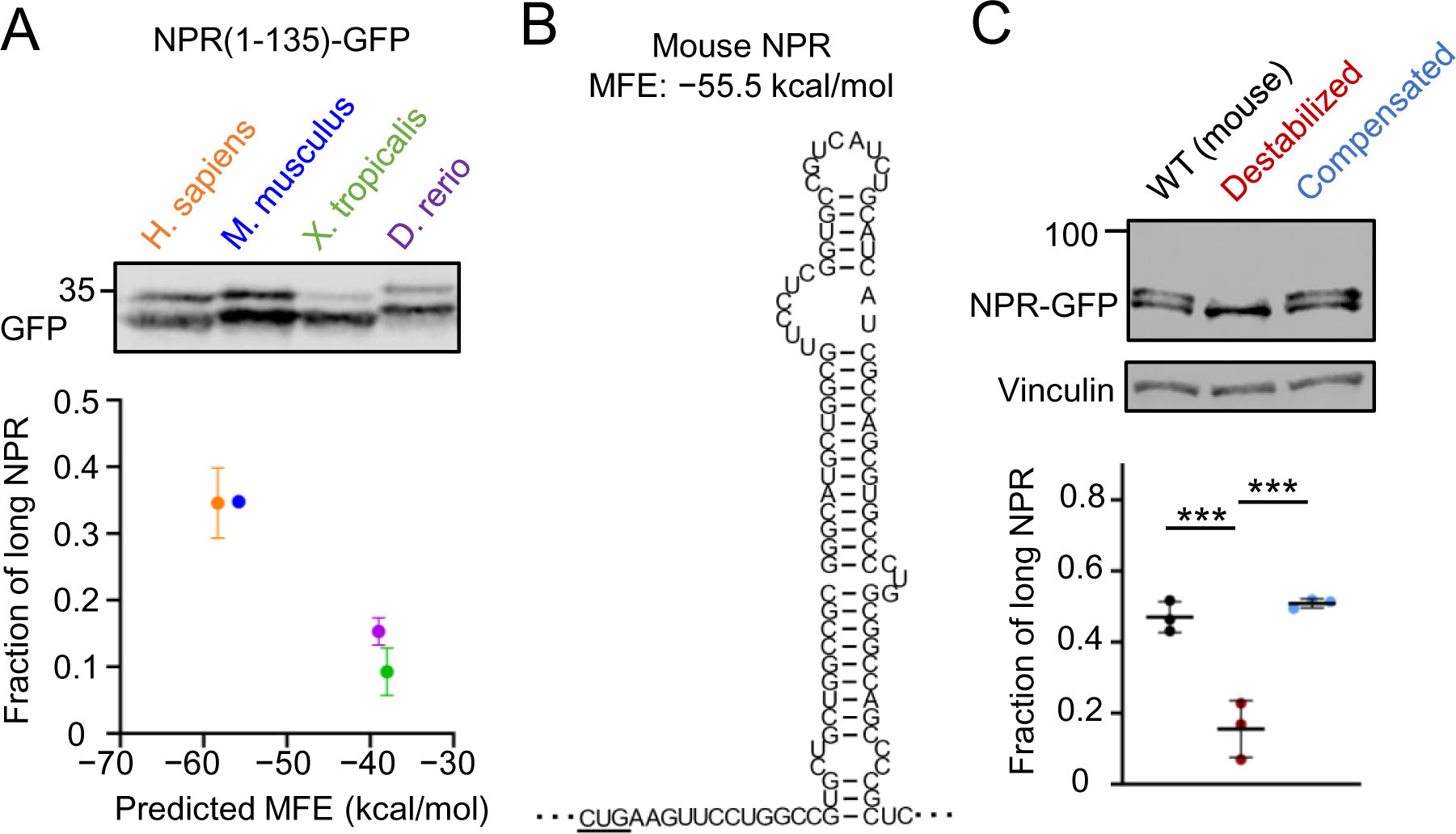
A stable RNA secondary structure promotes CUG initiation in mammals. (**A**) Proteoform ratios (*top*) and their correlation with the predicted MFEs (*bottom*) of four NPR orthologs. Data represent mean ± SD. n=3 independent experiments. (**B**) Predicted secondary structure of the first 100 nt of mouse NPTXR coding sequence. The CUG TIS and minimum free energy (MFE) are shown. (**C**) Proteoform ratios of wild-type (WT), destabilized, and compensatory mutant NPR cDNAs expressed in N2A cells. Data represent mean ± SD. n=3 independent experiments. ***, *P*<0.001, Student’s t tests.

Previous studies have shown that downstream RNA secondary structures can enhance initiation efficiency at both near-cognate TISs as well as AUG TISs within weak Kozak contexts (Kozak, 1990; Wang et al., 2022). Interestingly, *in silico* analysis (Gruber et al., 2008) predicted the presence of stable RNA stem-loop structures located 10-12 nucleotides downstream of the CUG start codon in human and mouse NPR mRNAs, with predicted minimum free energies (MFE) of−58.3 and −55.5 kcal/mol, respectively (**Figure 2A,B)**. In comparison, predicted secondary structures in frog and zebrafish NPR mRNAs were less stable (MFE of −38.0 and −39.1 kcal/mol) (**Figure 2A**). To test the functional significance of the predicted RNA structure, we first introduced synonymous mutations to the left arm to destabilize the stem region (MFE = −39.3 kcal/mol, **Figure S2**), which substantially reduced CUG initiation efficiency (**Figure 2C**). To further establish the causal role of the structure, we restored the base pairs by introducing compensatory mutations to the right arm of the stem (MFE = −58.5 kcal/mol). Expression of long NPR was fully restored by compensatory mutations (**Figure 2C**), indicating that the NPR mRNA secondary structure is both required and sufficient to enhance initiation at the CUG TIS in mammals.

### Alternative AUG initiation converts NPR to a secreted proteoform

Similar to other type II single-pass membrane proteins, the N-terminal transmembrane domain of CUG-initiated NPR (amino acids 3-23) presumably acts as a signal anchor to target the ribosome/mRNA complex to the ER surface. Therefore, N-terminal truncation caused by downstream AUG initiation may affect the site of translation and/or protein localization. To test whether the truncated N-terminus could still function as a signal anchor, we fused the N-terminal sequences from either the long or short NPR to GFP and monitored both cell-surface and intracellular expression. As anticipated, the intact signal anchor from long NPR was sufficient to direct GFP expression onto the outer surface in both HEK293T cells (**Figure 3B)** and primary hippocampal neurons (**Figure 3C**). In contrast, the truncated N-terminal sequence from short NPR resulted in no detectable GFP signals on the cell surface (**Figure 3B, C)**, suggesting that the truncated N-terminal sequence can no longer function as a signal anchor.

**Figure 3.**
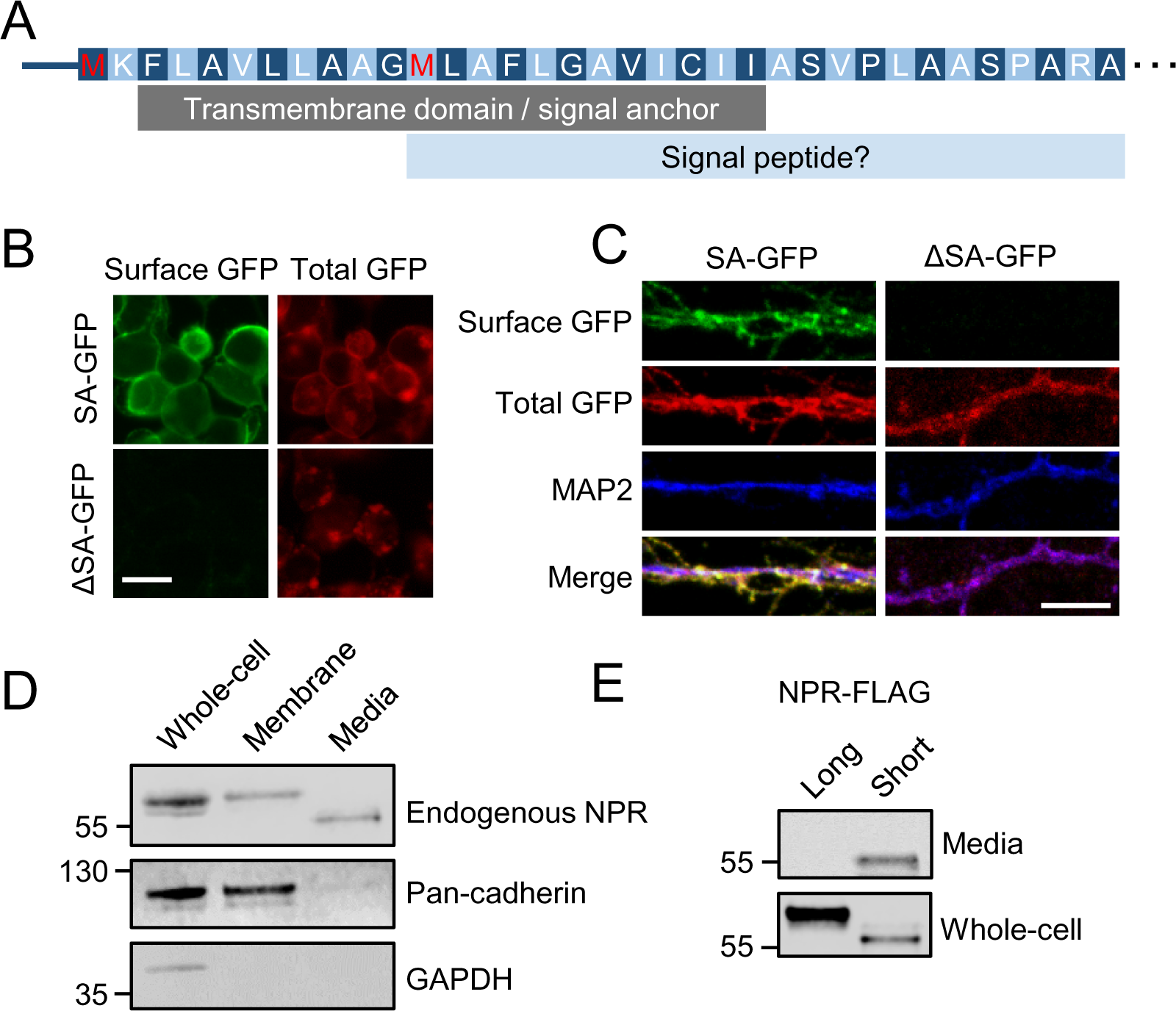
Alternative AUG initiation converts NPR to a secreted proteoform. (**A**) N-terminal sequence of NPR, showing the transmembrane domain/signal anchor (gray) and predicted signal peptide (blue). (**B**) Surface and total signals in HEK293T cells expressing GFP with either intact (SA) or truncated (ΔSA) N-terminus of NPR. Scale bar, 20 µm. (**C**) Surface and total GFP signals in hippocampal neurons expressing either SA- or ΔSA-GFP. Scale bar, 5 µm. (**D**) Endogenous NPR expression in cell lysates, membrane fractions, and culture medium of primary cortical neurons. Pan-cadherin and GAPDH are shown as membrane and cytosolic markers. (**E**) Expression of long and short NPR cDNAs in culture medium and whole-cell lysates of primary cortical neurons.

Instead, the truncated N-terminal sequence of short NPR was computationally predicted with high confidence as a cleavable signal peptide (Kall et al., 2004; 2007) (**Figure 3A, Figure S3A**), suggesting that short NPR may be directly secreted from cells. Indeed, not only was endogenous NPR detected in the culture media, but it also showed lower molecular weight than the NPR proteoform in the membrane fraction from primary cortical neurons (**Figure 3D**). Furthermore, expression of the long proteoform alone by substituting CUG with AUG did not generate detectable NPR in the media **(Figure 3E)**, arguing against the previously shown cleavage of long NPR by tumor necrosis factor-α converting enzyme (TACE) (Cho *et al*., 2008) as the main source of NPR secretion. Conversely, expression of short NPR alone (N-terminal deletion) resulted in robust secretion (**Figure 3E**), validating the *in silico* prediction that downstream TIS usage transformed the signal anchor to a cleavable signal peptide and thus converted transmembrane NPR into a secreted factor similar to NP1 and NP2.

### Secreted NPR enhances synaptic clustering of AMPA receptors *in vitro*

Both NP1 and NP2 have been shown to be regulated by neuronal activity (Schaukowitch et al., 2017; Tsui et al., 1996). To test whether the NPR proteoform ratio may also be regulated by neuronal activity, we treated primary cortical neurons with bicuculline, a GABA_A_ receptor antagonist, and monitored both intracellular and extracellular NPR proteoform expression. While long and short NPR in cells were largely unaffected, secreted NPR level was increased in the media after stimulation (**Figure 4A**), suggesting that either TIS usage or proteoform property may be sensitive to neuronal activity. To distinguish between these two possibilities, we performed TIS mapping by ribosome footprint profiling after treating neurons with harringtonine and puromycin to enrich ribosomes specifically at TISs (Gao et al., 2015). As expected, the inferred P sites of most ribosome footprints mapped to AUG, followed by near-cognate TISs (**Figure S4A**), confirming the enrichment of initiating ribosomes. Consistent with the observed increase in secreted NPR, ribosome footprints with their inferred P sites mapping to the downstream AUG were also increased relative to CUG after stimulation (**Figure 4B**), suggesting that alternative TIS usage in NPR mRNA was regulated by neuronal activity.

**Figure 4.**
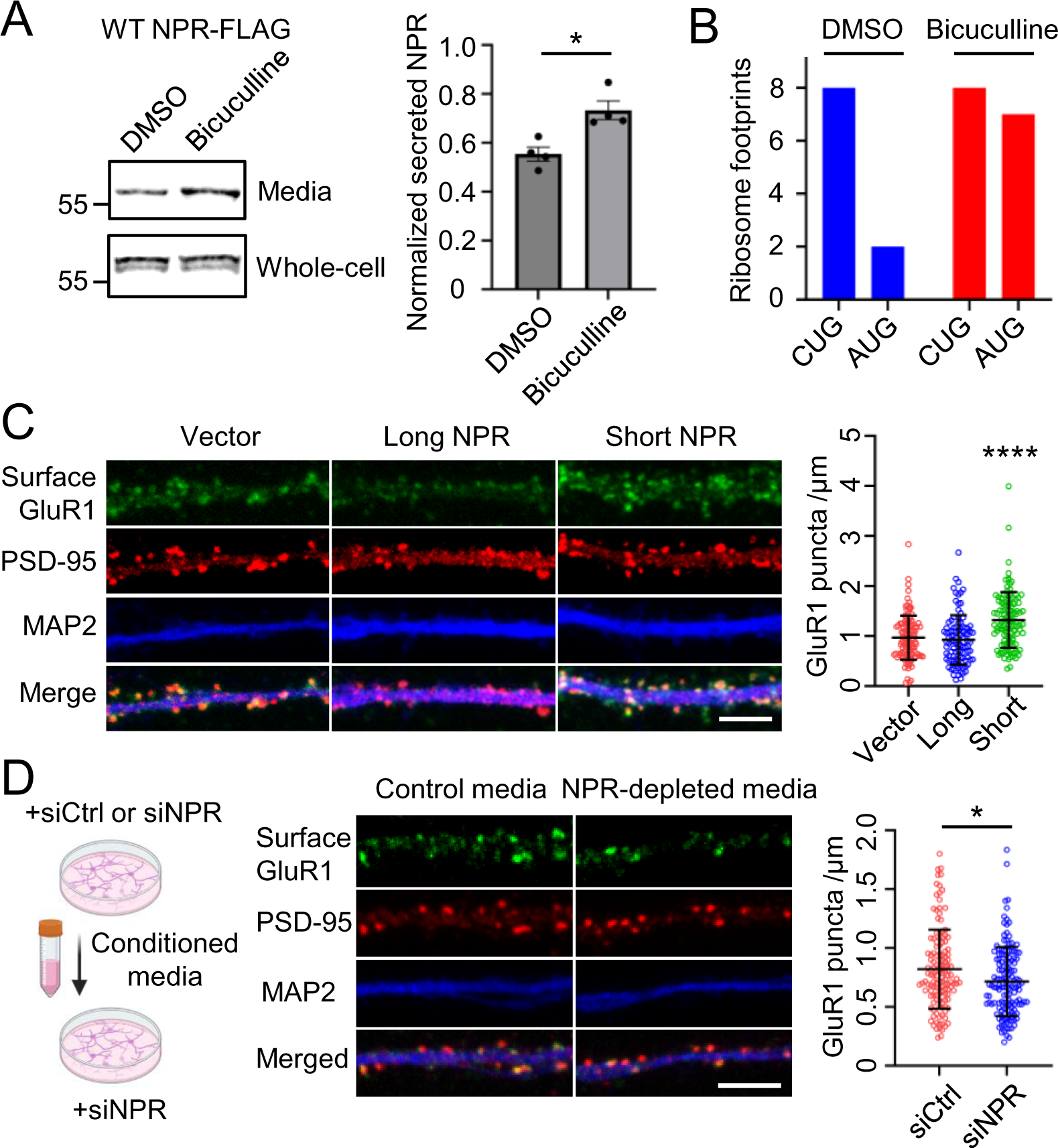
Secreted NPR enhances synaptic clustering of AMPA receptors *in vitro*. (**A**) Left: Western blot analysis of NPR secretion after DMSO or bicuculine (40 µM, 8 hour) treatment. Right: Quantification of secreted NPR levels normalized to long NPR levels in cells. (**B**) Numbers of ribosome-protected footprints with their inferred P-sites mapping to either CUG or AUG start codon after DMSO/bicuculine and harringtonine/puromycin treatment. (**C**) Surface GluR1, PSD-95 and MAP2 staining (*left*) in cultured hippocampal neurons expressing either FLAG (vector), long, or short NPR cDNA. Scale bar, 5 µm. Quantification of surface GluR1 puncta density in three independent experiments. ****, *P*<0.0001, Kruskal-Wallis tests comparing between short NPR and either vector or long NPR groups. (**D**) Left: Schematic illustration of the experimental procedure testing the effects of secreted NPRs in the media. Middle: Surface GluR1, PSD-95 and MAP2 staining in NPR-depleted recipient hippocampal neurons treated with conditioned medium with or without NPR depletion. Scale bar, 5 µm. Right: Quantification of surface GluR1 puncta density in three independent experiments. *, *P*<0.05, Mann-Whitney test.

The differential regulation of long and short NPRs by neuronal activity suggested that these two proteoforms might be functionally distinct. To test whether long and short NPRs may have differential impact on post-synaptic AMPA receptor clustering, we replaced endogenous NPR with individual proteoforms in primary hippocampal neurons by first knocking down endogenous NPR expression with a 3ʹ UTR-targeting siRNA (**Figure S4B**) and then overexpressing siRNA-resistant long or short NPR cDNAs. We monitored the synaptic clustering of AMPA receptors in primary hippocampal neurons by labeling them with a GluR1 antibody under non-permeabilized conditions and compared neurons expressing each individual NPR proteoform. Long NPR expression alone did not increase the density of surface GluR1 puncta (**Figure 4C**) but reduced their intensity (**Figure S4C**), possibly due to dominant negative effects caused by overexpressed long NPRs mislocalized outside of the pre-synaptic terminals. In contrast, short NPR expression alone significantly increased both the density (**Figure 4C**) and intensity (**Figure S4C**) of surface GluR1 puncta. These results suggest that long and short NPRs are functionally distinct, and that short NPR can function independently of its transmembrane counterpart in promoting AMPA receptor clustering.

Considering that short NPR was secreted into the media, we asked whether the effect of short NPR on AMPA receptor clustering required direct cell-cell contact (i.e., synaptic connections), or if it could be in part mediated by diffusion. We first collected control and secreted NPR-depleted conditioned media from primary neurons treated with either control or NPR-targeting siRNAs, respectively, then added these conditioned media to new batches of NPR-depleted neurons (**Figure 4D**). Neurons treated with control media showed higher surface GluR1 puncta density than those treated with NPR-depleted media (**Figure 4D**), indicating that short NPR could promote AMPA receptor clustering at least in part through secretion and diffusion, independent of direct cell-cell contact.

### Change in NPR proteoform ratio reduces GluR4 in PV neurons and causes behavioral changes *in vivo*

Previous studies have shown that neuronal pentraxins play crucial roles in the formation and organization of excitatory synapses on PV neurons (Chang et al., 2010; Pelkey *et al*., 2015). While knocking out both NP2 and NPR in mice causes a nearly complete loss of GluR4-containing AMPA receptors in PV neurons in the hippocampus (Pelkey *et al*., 2015), conventional gene knockout studies cannot address proteoform-specific contributions. To specifically alter proteoform ratio without changing the overall NPR abundance, we designed a proteoform-specific targeting strategy based on our finding that alternative initiation at downstream AUG required leaky scanning at the CUG TIS (**Figure 1D**). Using CRISPR/Cas9-based homologous recombination, we generated knock-in mice carrying one or both *Nptxr* alleles with the CTG TIS replaced with an ATG. Homozygous *Nptxr*^AUG^*/Nptxr*^AUG^ (hereafter referred to as AUG KI) mice were born at expected Mendelian ratios and did not display gross physical or behavioral abnormalities. Consistent with the expectation that substituting the CUG TIS with an AUG should effectively block leaking scanning, AUG KI mice expressed predominantly long NPR (**Figure 5A**), whereas the total NPR level remained unchanged. Importantly, immunohistochemistry showed that GluR4 levels were significantly reduced in PV neurons in the CA1 region of the hippocampus (**Figure 5B**), suggesting that the NPR proteoform diversity indeed contributed to AMPA receptor expression in these interneurons. Consistent with the reduced excitation of PV neurons, perineuronal net density measured by Wisteria floribunda agglutinin (WFA) labeling was also reduced in AUG KI mice (**Figure S5A**).

**Figure 5.**
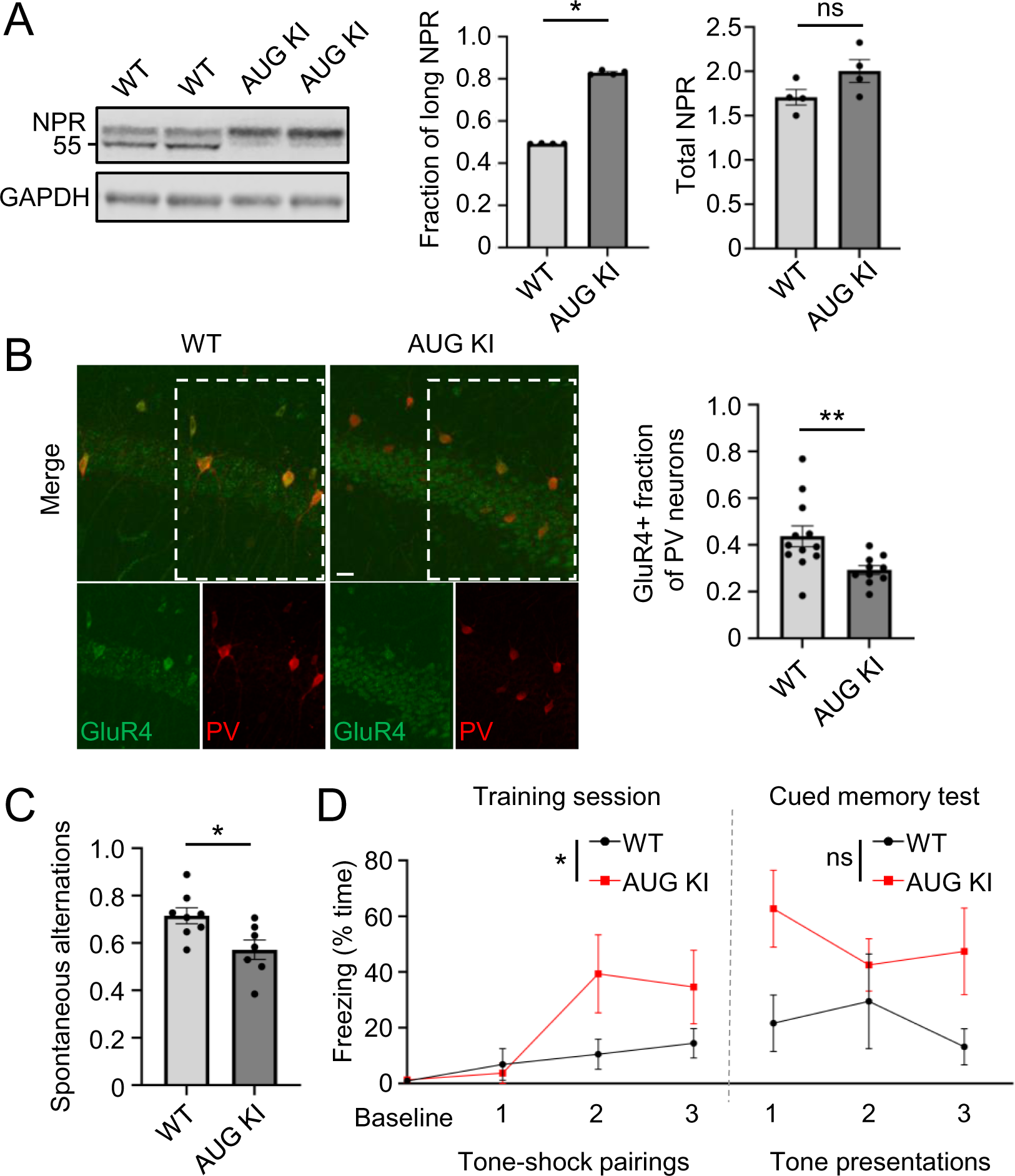
Change in NPR proteoform ratio reduces GluR4 in PV neurons and causes behavioral changes *in vivo*. (**A**) Left: Endogenous NPR expression in mouse forebrain lysates from wild-type (WT) and homozygous AUG knock-in (KI). Right: Long NPR fractions and total NPR abundance were quantified in n=4 animals per genotype. (**B**) Representative images (left) and quantification (right) of GluR4 and PV immunohistochemistry in CA1 regions of P30 WT and AUG KI mice (scale bar, 20 µm). n=3 per genotype. **, *P*<0.01, Mann-Whitney test. (**C)** Frequencies of spontaneous alternations in a Y-maze. *, *P*<0.05, Mann-Whitney test. (**D**) Freezing responses to tone preceding shock during training session (left) and to tone presentation during Cued Memory Test (right). n=5 animals per genotype. *, P<0.05 interaction; ns, P>0.05, 2-way Repeated Measures ANOVA.

PV neurons are broadly involved in cognitive functions and behaviors, with a particularly important role in working memory (Murray et al., 2015). We therefore tested AUG KI mice and their WT littermates in a number of behavioral paradigms that measure baseline function and different forms of short-and long-term memory. While WT and AUG KI mice showed similar ambulatory locomotor activity (**Figure S5B**), AUG KI mice made fewer spontaneous alternations in a Y-maze compared to WT littermates (**Figure 5C**), suggesting that AUG KI mice exhibited impaired working memory. In a separate paradigm, AUG KI mice displayed increased within-session freezing to a shock-paired tone after the first tone-shock pairing, suggesting that loss of NPR proteoform diversity increased the sensitivity to a stressful stimulus and enhanced short-term associative fear learning (**Figure 5D**). Compared to the immediate enhancement, the difference in freezing in response to the tone alone 48 hours later was diminished, suggesting that long-term fear memory was less affected (**Figure 5D**). The increase in short-term tone-fear reactivity and the reduction of spontaneous alternations in a Y-maze may be due to a differential role for PV neuron activity in fear learning (Yau et al., 2021) versus working memory (Murray *et al*., 2015). Taken together, these results indicated that a large change in NPR proteoform ratio caused both cellular and behavioral changes *in vivo*, supporting the notion that NPR proteoform diversity is necessary for appropriate development of PV neurons as well as cognitive behaviors that rely on intact PV neuron function.

### N-terminal extension by upstream alternative TISs converts secreted C1QLs to transmembrane proteoforms

Having observed the signal anchor-to-signal peptide conversion through N-terminal truncation of NPR, we reasoned that the reverse conversion might also occur when alternative translation initiation occurred at an upstream alternative TIS. Among the candidate proteins previously predicted by sequence analysis to have conserved N-terminal extensions (Ivanov *et al*., 2011) are the complement component 1, q subcomponent-like proteins (C1QLs), which have been shown to regulate the formation and maintenance of excitatory synapses (Martinelli et al., 2016; Matsuda et al., 2016). In contrast to NPR, C1QLs have been identified as secreted synaptic organizers indirectly anchored at the presynaptic terminal through one or more membrane proteins such as neurexins (Matsuda *et al*., 2016) and bind to several postsynaptic partners including the cell-adhesion G protein-coupled receptor BAI3 (Bolliger et al., 2011) and kainate receptors (Matsuda *et al*., 2016). Intriguingly, all four C1QL paralogs have PhyloCSF candidate coding regions (Lin et al., 2011) with high ratios of synonymous to non-synonymous mutations indicative of conserved protein-coding functions immediately upstream of their canonical AUG TISs, each proceeded by a conserved AUU codon within an ideal Kozak context (**Figure 6A**).

**Figure 6.**
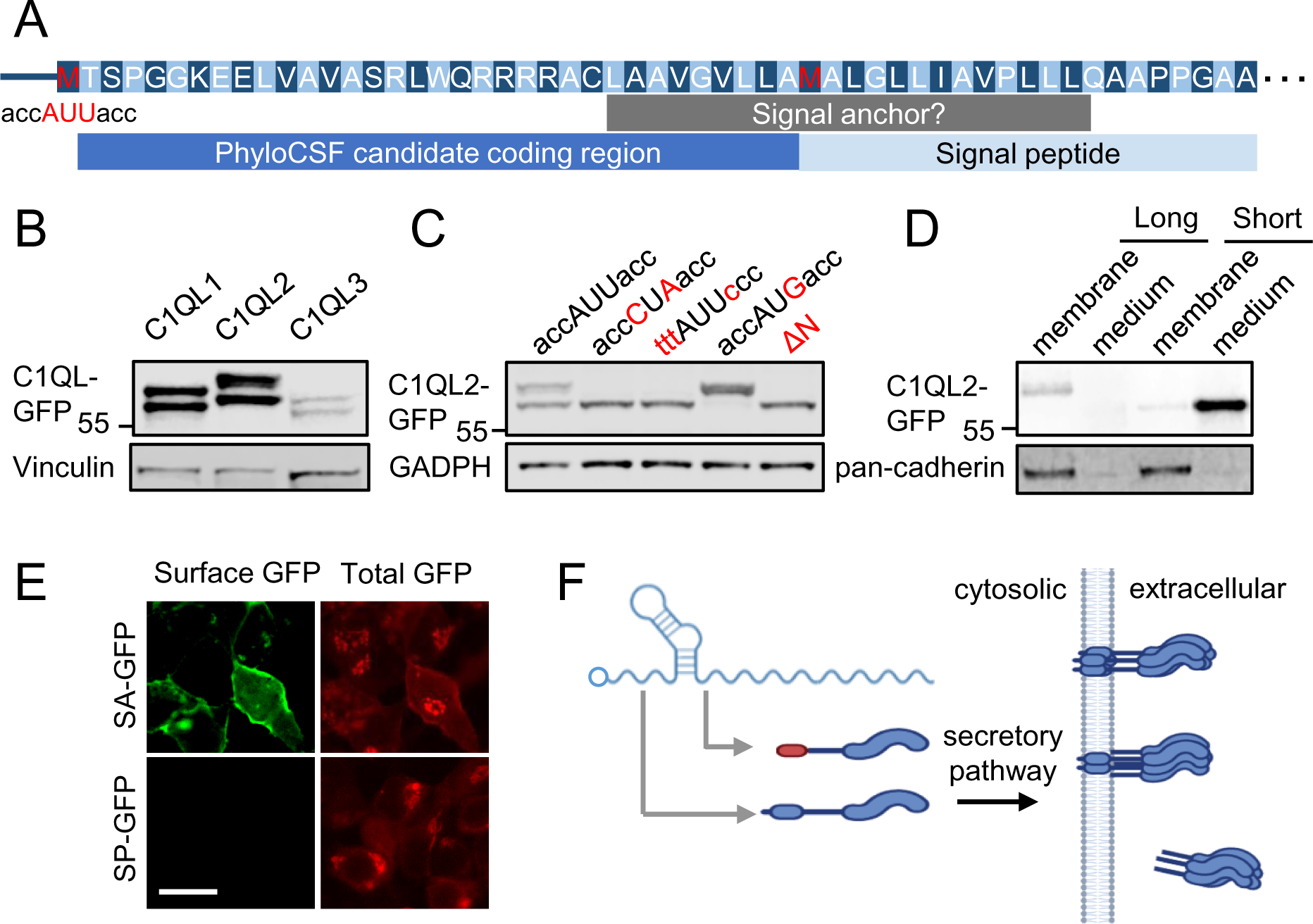
N-terminal extension by upstream alternative TISs converts secreted C1QLs to transmembrane proteoforms. (**A**) N-terminal sequence of C1QL2, showing the putative upstream AUU TIS (red), PhyloCSF candidate coding region, predicted SA and SP sequences. (**B**) Ectopic expression of WT C1QL1, C1QL2, and C1QL3 cDNAs in HEK293T cells. (**C**) Expression of mutant C1QL2 cDNAs in HEK293T cells. (**D**) Expression of long and short C1QL2 in membrane fractions and culture medium of primary cortical neurons. (**E**) HEK293T cells expressing GFP with the putative SA or SP sequences from long and short C1QL2, respectively. Scale bar, 20 µm. (**F**) Diagram illustrating the role of alternative translation initiation enabling mRNAs to dually encode transmembrane and secreted proteoforms.

Consistent with a previous study detecting multiple C1QL2/3 protein species by immunoblotting (Matsuda *et al*., 2016), ectopically expressed C1QL1/2/3 along with their native 5ʹ UTRs also each produced two proteoforms (**Figure 6B**). The apparent molecular weights of C1QL2 proteoforms were similar to those of two mutants in which either the AUU codon was substituted by an AUG or the putative N-terminal extension was deleted (ΔN) (**Figure 6C**). Either substituting the AUU with CUA or changing the AUU-flanking sequence to an anti-Kozak context eliminated the expression of long C1QL2 proteoform (**Figure 6C**), confirming the AUU codon as the upstream alternative TIS. Similar to the CUG-downstream sequence in NPR, AUU-downstream sequences in C1QL1-4 were also predicted to form stable RNA secondary structures (**Figure S6A**).

Consistent with C1QLs being known as secreted proteins, all four N-terminal sequences of canonical AUG-initiated short C1QL proteoforms were predicted as cleavable signal peptides (**Figure S6B**). In contrast, all four AUU-initiated long C1QL proteoforms were predicted to have N-terminal transmembrane domains (**Figure S6B**), suggesting potential conversions from signal peptides to signal anchors. To test these predictions, we expressed each of the two C1QL2 proteoforms in primary cortical neurons. While short C1QL2 was mostly secreted into the media as expected, long C1QL2 was predominantly membrane-bound (**Figure 6D**). Furthermore, the extended N-terminal sequence of long C1QL2, but not the canonical N-terminal sequence of short C1QL2, was sufficient to direct the cell-surface expression of GFP (**Figure 6E**), consistent with the *in silico* prediction of a signal peptide-to-signal anchor conversion. These results, together with those from NPR, indicated an unexpected level of plasticity of N-terminal signal sequences tuned by alternative start codon usage, which enabled dual encoding of secreted and transmembrane proteoforms (**Figure 6F**).

### Prediction of additional alternative TIS-mediated proteoform conversions

To explore the broader scope of this mechanism, we asked whether alternative TIS-mediated signal anchor/signal peptide conversions could be found in other mRNAs beyond those encoding synaptic organizers. Using a list of alternative TISs identified by previous ribosome profiling studies in human (Chen *et al*., 2020) and mouse cells (Ingolia *et al*., 2011), we applied Phobius (Kall *et al*., 2004; 2007) to predict signal peptides and transmembrane domains for each N-terminal variant and their annotated protein sequences. This analysis identified 35 secreted proteins with cleavable signal peptides that were predicted to acquire a transmembrane domain (putative signal anchor) through upstream TIS-mediated N-terminal extension, as well as 22 proteins with single predicted transmembrane domains that were changed to cleavable SPs upon N-terminal truncation (**Supplementary Table 1**). These putative N-terminal proteoforms included an additional C1q-related factor (C1QTNF1) as well as a wide variety of functional categories including extracellular matrix components (e.g., laminin), signaling molecules (e.g., WNT3/5A, gremlin), and cell adhesion molecules. In addition, this analysis also predicted N-terminal truncation events that removed the N-terminal signal sequences, possibly resulting in the retention of these ER-targeted protein within the cytoplasm, as well as the acquisition of a putative N-terminal signal sequences by cytoplasmic proteins through N-terminal extension.

## DISCUSSION

In this study, we show that a conserved CUG TIS in NPR mRNA enables alternative translation initiation at the downstream AUG TIS, yielding an N-terminal truncated proteoform with distinct subcellular localization and synaptic functions. By generating a mouse model with an altered NPR proteoform ratio, but not the overall NPR abundance, we found that the balance between the expression levels of long and short NPR proteoforms plays a role in regulating GluR4 and excitation levels in PV interneurons in the hippocampus. Concomitantly, the loss of NPR proteoform diversity affected cognitive functions, with mice expressing predominantly long NPR exhibiting sensitized short-term associative fear learning and impaired working memory measured by spontaneous alternations in a Y maze.

In contrast to the widely accepted notion that nearly all multi-exon genes produce alternatively spliced mRNA isoforms, the conventional view has been that each of the vast majority of mRNAs encodes a single protein. This view has been challenged most notably by ribosome profiling studies, particularly those that specifically enrich post-initiation ribosomes to globally map TISs across the transcriptome (Gao *et al*., 2015; Ingolia *et al*., 2011). Instead of finding one TIS per mRNA, these studies have repeatedly shown that alternative TIS usage is much more prevalent than previously thought, with more than half of the detected transcripts having more than one TIS. Using these expanded TIS annotations, we searched for additional N-terminal signal sequence switching events analogous to NPR and the C1QL proteins. Indeed, we were able to identify dozens of other mRNAs encoding distinctly localized proteoforms, hinting at a broad impact of alternative TIS usage on protein localization. However, ribosome profiling-based TIS mapping can conceivably cause false positives. For example, to eliminate the footprints protected by actively translating ribosomes, these experiments typically involve treating cells with translation inhibitors over an extended period of time, thereby allowing translating ribosome to run off. During this extended treatment, new initiation events might accumulate at positions that may or may not represent physiological TIS usage. To partly address this caveat, QTI-seq enriched initiating ribosomes after cell lysis, thereby minimizing new rounds of initiation (Gao *et al*., 2015). Furthermore, puromycin was added to induce the release of the nascent polypeptide chain and the translating ribosome, which facilitated the depletion of elongating ribosomes. Nonetheless, most alternative TISs will require validation by orthogonal approaches. In the cases of NPR and C1QLs, their alternative TIS usage was confirmed by both evolutionary conservation (e.g., PhyloCSF) and protein-level detection. Additionally, high-throughput validation of bifunctional mRNA candidates can also be achieved through mass spectrometry-based proteomics (Van Damme *et al*., 2014).

The biogenesis of transmembrane and secreted proteins heavily relies on their N-terminal signal sequences. Type II single-pass transmembrane proteins use an N-terminal signal anchor that also functions as a transmembrane domain, whereas secreted proteins have cleavable signal peptides. While both types of signal sequences induce ER targeting of the mRNA-ribosome-nascent peptide chain complex through the SRP complex, these proteins enter unique downstream processing mechanisms, possibly at different subdomains of the ER (Lak et al., 2021). The processing of signal anchors versus signal peptides requires distinct ER-resident factors capable of detecting the differences between the two types of signals. For instance, the nascent polypeptide chain of a type II transmembrane protein is inserted into the ER membrane head-first and undergoes a signal anchor inversion within the ER-Sec61α complex, bypassing signal peptidase-dependent cleavage of signal peptides of secreted proteins. In the context of bifunctional mRNAs that produce both transmembrane and secreted proteoforms, whether TIS selection can alter their localization to different ER subdomains remains to be tested.

A related question with regards to mRNAs with multiple TISs is whether all TISs are used in each mRNA copy. Translation of a single copy of NPR mRNA, for example, may be initiated at either CUG, AUG, or both TISs. Single-molecule imaging-based analysis of mRNA translation may elucidate the mechanistic details underlying ER subdomain-specific control of alternative TIS selection (Wu et al., 2016). Considering the distinct morphology of neurons characterized a high degree of compartmentalization, TIS selection of many neuronal mRNAs may also depend on the subcellular localization (Wu *et al*., 2016). Furthermore, the subcellular localization of NPR mRNA and/or its local translation might also underlie the observed activity-dependent regulation of TIS usage.

In addition to potential subcellular compartment-specific TIS usage, our analysis of NPR cDNA expression across different cell types indicated a high long-to-short NPR proteoform ratio in neurons. While mouse NPR mRNA expressed 35% long proteoform in rabbit reticulocyte lysate and 47% in N2A cells, both endogenous and ectopically expressed NPR mRNA produced 62% and 83% long proteoform in primary neurons, respectively, suggesting that one or more factors that regulate alternative TIS usage, such as eIF1, eIF1A, and eIF5, may be differentially expressed between cell types. Considering that efficient initiation at the CUG TIS requires a highly stable downstream stem-loop structure, the folding equilibrium of the structure as well as its unwinding during ribosome scanning may also be regulated by cellular factors differentially expressed between cell types. Furthermore, the usage of non-AUG TISs is known to be regulated by cell states, with several stress-responsive mRNAs (e.g., GADD45G) reported to be translated at alternative TISs under cellular starvation (Gao *et al*., 2015), raising the possibility that the proteostasis collapse in neurodegenerative disorders may be in part due to widespread changes in alternative TIS usage in neuron under stress.

While our study on synaptic organizers primarily focused on the alternative TIS-mediated plasticity of N-terminal signal sequences, alternative translation can conceivably impact protein functions in diverse ways. For instance, alternative TIS may add or subtract critical residues for post-translational modification or protein-protein interactions potentially impactful on protein folding and catalytic functions. In addition, alternative TISs of upstream ORF (uORF) and out-of-frame TISs can regulate the amount of functional protein output. Therefore, our findings on alternative translation initiation, together with other modes of translation recoding mechanisms such as ribosomal frameshifting and stop-codon read-throughput, hint at a vastly underexplored contribution of alternative translation in diversifying the functional output of the transcriptome.

### Limitations of the study

In order to alter the relative proteoform ratio while keeping the overall NPR abundance constant, we generated a CTG-to-ATG KI mouse model to shift the relative ratio towards long NPR. As such, while the cognitive and synaptic phenotypes observed in these mice support the significance of the proper proteoform ratio, the underlying mechanism remains to be further investigated. In one scenario, these phenotypes may result from either the near-complete loss of short NPR and/or the overexpression of the long NPR. In a separate scenario, the proteoform ratio may represent the correct subunit stoichiometry critical for the formation of functional heteromeric complexes either between long and short NPRs or between NPR and NP1/NP2, as previously reported (Xu et al., 2003). Lastly, a previous study has shown that (long) NPR can undergo TACE-mediated cleavage (Cho *et al*., 2008), which in turn regulates AMPA receptor clustering. Although we did not observe substantial level of long NPR cleavage in primary neurons, changes in subunit stoichiometry could conceivably affect the efficiency of TACE cleavage. Future in vitro and in vivo studies with additional proteoform-specific perturbations may help disentangle these possibilities.

## METHODS

### Animals

Animal care and housing were provided by the Yale Animal Resource Center (YARC). The animals were maintained in a 12-hour light/dark cycle with ad libitum access to food and water. All experimental protocols involving animals were reviewed and approved by the Institutional Animal Care and Use Committee (IACUC) at Yale University (Protocol #2021-20207). AUG KI mice carrying one or both *Nptxr* alleles with the CTG-to-ATG codon replacement were generated by Yale Genome Editing Center using CRISPR/Cas9-based homologous recombination. Sequencing-validated founder animals were backcrossed to WT C57BL6/J animals to establish a homogenous genetic background. The mutant line was subsequently maintained through het-to-het breeding. For behavioral and immunohistochemical experiments, homozygous AUG KI mice and their WT littermates were used.

### In vitro translation

In vitro translation was performed using a rabbit reticulocyte lysate system (Promega, L4960). Capped and polyadenylated reporter mRNAs containing the first 135 nucleotides of NPR coding sequences (human, mouse, frog, or zebrafish), followed by the GFP sequence were generated by using the HiScribe T7 ARCA mRNA Kit (New England Biolabs, E2060). 1 µg mRNA was first denatured at 65°C for 3 minutes, and then added to the rabbit reticulocyte lysate reaction mix. The translation reactions were incubated for 90 minutes at 37 °C for human, 36.6°C for mouse, 25°C for frog, and 28.5°C for zebrafish reporter mRNA. The translation products were analyzed by immunoblotting.

### AAV preparation and transduction

HEK293T cells were cultured in DMEM supplemented with 10% fetal bovine serum (FBS). Cells were transfected with pAAV-hSyn plasmid containing the cDNA of interest, along with the AAV-DJ and pHelper constructs, to produce AAV in a T175 flask for 48 h. AAV-producing cells were detached by incubation with 0.5 M EDTA (pH 8.0) at room temperature for 10 min. Harvested cells were then centrifuged at 2,000 g for 10 min at 4°C to remove the supernatant. Viruses were extracted and purified by using an AAV extraction kit (Takara).

### Primary neuronal culture and AAV transduction

Pregnant C57BL/6J mice (Charles River) at 18 days gestation were anesthetized with isoflurane for 5 min in a chamber and subsequently sacrificed by cervical dislocation. E18 embryos were then extracted and transferred to a sterile dish containing HBSS. Brain regions of interest, such as cortices or hippocampi, were dissected, removing any surrounding tissue and meninges. The tissues were digested in 5 mL of trypsin-EDTA solution and incubated at 37°C for 10-15 min, followed by gentle tituration in DMEM supplemented with 10% FBS (Thermo Fisher). Neurons were pelleted and resuspended in Neurobasal medium supplemented with B27 Plus (Thermo Fisher, A3582801, 50X), L-Glutamax (Thermo Fisher, 35050061, 1%), 33 mM glucose, and 37.5 mM NaCl. Dissociated neurons were plated on coverslips or dishes coated with laminin and poly-ornithine at the desired density. The medium was changed every 3-5 days, and 5 µM AraC was added on day 1 in vitro (DIV1) for 48 hours to inhibit glial proliferation.

For NPR replacement experiments, neurons were treated with 1 µM siRNA (Dharmacon: A-046750-16-0050) targeting the 3ʹ UTR of NPR mRNA (5ʹ-CTTGCAAACTGAATTCCTA-3ʹ) at DIV2. NPR siRNA was supplemented at every medium change. Neurons were transduced with AAV for overexpression of WT or mutant NPR cDNAs simultaneously with siRNA treatment. Neurons were allowed to mature until DIV14-16 for further analyses.

### Subcellular fractionation

Cells were harvested in ice-cold subcellular fractionation buffer (20 mM HEPES, pH 7.4, 10 mM KCl, 2 mM MgCl_2_, 1 mM EDTA, 1 mM EGTA, 1 µM DTT, supplemented with a cocktail of protease inhibitors) and incubated on ice for 15 min before being passed through a 26-gauge needle 10 times and kept on ice for additional 20 min. The cell lysates were first centrifuged at 720 g for 5 min at 4°C to pellet the nuclei, then further centrifuged at 10,000 g for 5 min at 4°C to pellet the mitochondria. The remaining supernatant was ultracentrifuged at 100,000 g for 1 h at 4°C. The pellet containing the membrane fraction was washed in 2M KCl for 1 h at 4°C to remove peripheral membrane proteins and subsequently ultracentrifuged for another 1 h at 4°C. The membrane pellet was finally resuspended in TBS containing 0.1% SDS.

### Synaptosome preparation

Freshly harvested brain regions from adult mice were suspended in ice cold Buffer A (320 mM sucrose, 10 mM HEPES pH 7.4, 1 µM DTT, supplemented with a cocktail of protease inhibitors). Tissues were homogenized using a glass Dounce tissue grinder (KIMBLE). Homogenates were centrifuged at 800 g for 10 min at 4°C. The supernatant was further centrifuged at 9,000 g for 15 min at 4°C. The pellet was resuspended and washed in ice-cold Buffer A and centrifuged again at 9,000 g for 15 min at 4°C to obtain crude synaptosomes.

### Western blotting

Cultured cells were lysed in RIPA buffer on ice for 10 min. After 10 min centrifugation at 4°C, 20,000 g, whole-cell lysates were mixed with 4X LDS sample buffer (Thermo Fisher) and denatured at 95°C for 5 min. Samples were loaded on a 4-12% Bis-Tris-SDS-PAGE gel, run at 120 V for 2 h 30 min in MOPS buffer, and transferred onto a nitrocellulose membrane (Bio-Rad) at 15V for 55 min. After 1 h blocking with 5% nonfat dry milk in TBS-T, the membrane was incubated with primary antibodies against protein of interest (NPR, Santa Cruz, sc-39008, 1:100; GFP, Aveslab, GFP-1020, 1:5,000; FLAG, Sigma, F1804, 1:1,000; FLAG, Rockland, 600-401-383, 1:1,000; GAPDH, Sigma, G9545, 1:2,000; Pan-Cadherin, Sigma, C1821, 1:1,000; Vinculin, Sigma, V9264, 1:5,000) diluted in 5% milk/TBS-T, placed on a shaker overnight at 4°C. After incubation, membranes were washed three times with TBS-T, followed by incubation with IR680-or IR800-conjugated secondary antibodies (Li-Cor, 1:5,000) diluted in 5% milk/TBS-T at room temperature for 1 h. Subsequently, membranes were washed with TBS-T and imaged using Odyssey XF system (Li-Cor, 2800).

### Immunoprecipitation

For immunoprecipitation of endogenous NPR, culture medium of primary cortical neurons was first centrifuged at 1,000 g for 5 min at 4°C to remove cell debris, and then incubated with an NPR antibody (Santa Cruz, sc-39008, 1:50) overnight at 4°C in a mini-rotator. Samples were then incubated with pre-washed sheep anti-mouse antibody (M280)-conjugated Dynabeads (Thermo Fisher, 11201D) for 1 h at room temperature. Subsequently, the beads were washed three times with PBS supplemented with 0.1% bovine serum albumin and 2 mM EDTA, pH 7.4. Bound proteins were eluted by boiling at 95°C in SDS sample buffer for 10-15 min and were analyzed by western blotting.

For FLAG- and GFP-tagged proteins, culture medium was incubated with either anti-FLAG antibody (M2)-conjugated magnetic beads (Sigma, M8823) or GFP-Trap magnetic agarose affinity beads (Chromotek, gtma-20) overnight at 4°C in a mini-rotator. The beads were washed three times with the appropriate washing buffer (TBS for Anti-FLAG M2 Magnetic beads; 10 mM Tris/Cl pH 7.5, 150 mM NaCl, 0.5 mM EDTA for GFP-Trap magnetic agarose), followed by boiling in SDS sample buffer at 95°C for 10-15 min to elute the bound proteins. Samples were then analyzed by western blotting.

To remove glycans from proteins, cell lysates were first incubated with Deglycosylation Mix Buffer 2 (NEB) at 75°C for 10 min. After cooling down to room temperature, Protein Deglycosylation Mix II (NEB) was added and further incubated for 30 min. The mixture was then transferred to a shaker and incubated at 37°C for 1 h.

### Immunocytochemistry

For live-cell labeling of GluRs, DIV14-16 primary hippocampal neurons cultured on coverslips were first incubated with 1.0% bovine serum albumin in Tyrode buffer (15 mM D-(+)-Glucose, 108 mM NaCl, 5 mM KCl, 2 mM MgCl_2_, 2 mM CaCl_2_, 1 M HEPES, pH 7.4) for 5 min at room temperature. Cells were then incubated with GluR1 antibody (Sigma, ABN241, 1:100) in Tyrode buffer for 12 min at room temperature. Subsequently, cells were washed with PBS and fixed with 4% paraformaldehyde in PBS for 15 min at room temperature. Cells were permeabilized with 0.25% Triton-X in PBS for 5 min and then blocked in 10% normal goat serum in PBS for 1 h at room temperature. For staining of intracellular proteins of interest, such as PSD-95 (Sigma, MAB1596, 1:500 diluted) and MAP2 (Synaptic Systems, 188 006, 1:1000), coverslips were incubated with the primary antibody diluted in 10% normal goat serum overnight at 4°C. After washing with PBS, cells were incubated with secondary antibodies conjugated to Alexa 488, Alexa 555, and Alexa 647 (Thermo Fisher, 1:500 in 10% normal goat serum). After washing, cells were mounted on glass slides in ProLong Diamond Antifade Mountant (Thermo Fisher).

For surface labeling of GFP, cells were fixed with 4% paraformaldehyde in PBS for 10 min at room temperature. After washing with PBS, fixed cells were blocked in 10% normal goat serum in PBS for 1h at room temperature, and then incubated with GFP antibody (Thermo Fisher, A6455, 1:500) in 10% normal goat serum for 2 h at room temperature. After washing with PBS, cells were incubated with secondary antibodies diluted in 10% normal goal serum for 1 h at room temperature. Subsequent staining of intracellular proteins was performed as described above.

### Image acquisition and analysis

Z-stack images of primary neurons were taken with a Zeiss LSM 800 laser scanning confocal microscope with Airyscan. All quantitative measurements were performed with ImageJ imaging software (NIH). Before quantification, maximum intensity projections for each image were generated. For the quantification of either surface GluR1 or PSD-95 puncta, we restricted our analysis to those residing within or immediately adjacent to MAP2-positive secondary dendrites by masking. After setting minimum intensity threshold level for each fluorescence channel, both the number and mean intensity of puncta were measured by using the ‘analyze particles’ function. The puncta counts were normalized by dendritic length to calculate puncta density. All statistics were performed by using either GraphPad Prism or RStudio. We used Mann-Whitney U tests for all pairwise comparisons and Kruskal-Wallis tests for comparisons between multiple groups.

### Locomotor activity

All rodent behavioral analyses used approximately equal numbers of 10-month-old male littermates at the time of testing. Mice were placed in the testing room at least 30 min before each assay. All experiments and analyses were performed completely in a genotype-blind manner. Locomotor data was collected using an Accuscan Instruments behavioral monitoring system and Fusion software (Omnitech Electronics). Mice were temporarily single-housed on a 12 hr light-dark cycle with food and water ad libitum. Nesting material was removed to prevent obstruction of infrared beams in the locomotor monitoring system. Locomotion was monitored for 72 hr using 12 photocells placed 4 cm apart. Locomotor counts were monitored in 60 s blocks to obtain an “ambulatory activity count” consisting of the number of beam breaks recorded during a period of ambulatory activity. Mice were not disturbed during the testing period. Raw data from the Fusion software was then post-processed using a custom MATLAB script to extract the first 3 hr of data and re-organize into 15 min time bins.

### Fear conditioning

Mice were tested in four 6-min sessions (Habituation, Training, Contextual Memory Test, and Cued Memory Test) across 4 days modified from previously described protocols (Park et al., 2016). All protocols were administered using VideoFreeze software (Med Associates). For Habituation on Day 1, mice were placed in a lit plastic chamber (Med Associates) with a cardboard floor and some bedding from the mouse’s home cage (Context A). No shocks or cues were delivered. For Training on Day 2, mice were placed in the same plastic chamber with white plastic covering the walls, light, stainless steel grids for shock delivery (Context B), and almond extract underneath the grids to provide a specific odor. After a 160-s baseline, a tone (85 dB, 2.8 kHz) was presented for 20 s, with the last 2 s coinciding with a foot shock (0.5 mA). This same tone-shock pairing was provided a total of 3 times with 40-s inter-trial intervals and a 60-s final resting period. For the Contextual Memory Test on Day 3, mice were again placed in the chamber set up for Context B, but with orange extract underneath the grids to provide the odor. No shocks or cues were delivered. For the Cued Memory Test on Day 4, mice were placed in the chamber set up for Context A, and the sound cues were administered as described for Training, but without foot shocks. Mice were returned to their home cage after each session. Freezing behavior during each session was analyzed by the VideoFreeze software, and then post-processed using a custom MATLAB script to organize the data.

### Y-maze spontaneous alternation test

Mice were acclimated to the testing room and handling for at least 3 days prior to the experiment. For the test, the animal was first placed at the center of the Y maze with all three arms initially blocked from entry. After removing the barricades, the animal was allowed to freely explore the three arms for 5 minutes. An arm entry was recorded when all four limbs of the test animal are within the arm. For each trial, the total numbers of arm entries and alternations were counted. The fraction of spontaneous alternation was then calculated as:

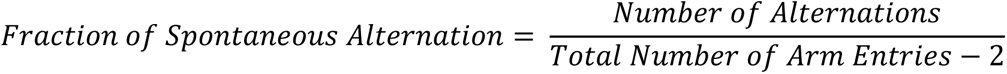

### Transmembrane domain and signal peptide prediction

Previously annotated human (Chen *et al*., 2020) and mouse (Ingolia *et al*., 2011) alternative N-terminal proteoforms based on ribosome profiling results, as well as their corresponding canonical proteoforms, were used to predict bifunctional signal sequences. Both transmembrane domains and cleavable signal peptides were simultaneous predicted by using Phobius (standalone version 1.01) (Kall *et al*., 2004; 2007). Signal anchor-to-signal peptide conversions were defined as N-terminal truncations that converted the canonical proteoform with a single transmembrane domain to a proteoform with a signal peptide and no predicted transmembrane domain. The opposite was defined as signal peptide-to-signal anchor conversions.

### TIS mapping by ribosome profiling

E16 primary cortical neurons were cultured in 10 cm dishes for 12 days before being treated with either DMSO or bicuculline (40 μM) for 10 hours. TIS mapping by ribosome profiling was performed by following the previously described QTI-seq method (Gao *et al*., 2015) with minor modifications. Neurons were washed twice briefly with ice-cold PBS supplemented with 5 μM harringtonine. 400 μl cold cell lysis buffer (20 mM HEPES, pH 7.4, 100 mM KCl, 5 mM, MgCl2, 500U/ml RNasin-Plus (Promega), 1x protease inhibitor cocktail (200X, Millipore), 5 μM harringtonine) was added directly to the cells, which were then scraped off of the dishes and transferred to 2 ml microcentrifuge tube containing Lysing Matrix-D (MP Biomedicals). Cells were lysed by vortexing 6 times (20 seconds each) with 40 s intervals on ice. After centrifugation for 10 min at 13,000 g at 4 °C. The supernatant was transferred into a new microcentrifuge tube and supplemented with 10 mM creatine phosphate, 0.1 mM spermidine, 40 μg/mL creatine phosphokinase, 0.8 mM ATP, and 25 μM puromycin, before incubation at 35 °C for 15 min. After incubation, RNA concentrations of the cell lysates were measured using Qubit RNA High Sensitivity Assay Kit (Thermo Fisher, Q32855). 10U RNase I (Thermo Fisher, EN0601) was added into cell lysate for every 100 μg RNA, and digestion was carried out at 4°C overnight with gentle rotation. The next day, RNase I digestion was stopped by adding Superase-In RNase inhibitor (Thermo Fisher, AM2696) at 1 U/μl. Digested cell lysate was loaded on top of 0.8 ml sucrose cushion (1 M sucrose, 20 mM HEPES, pH 7.4, 100 mM KCl, 5 mM, MgCl2, 5 μM harringtonine, 25 μM of puromycin) in a 1.5 ml ultracentrifuge tube (Beckman Coulter, 343778) and centrifuged for 1.5 h at 75,000 rpm at 4°C using a TLA-120.2 rotor (Beckman Coulter). After carefully removing the supernatant, the ribosome pellet was solubilized in TRIzol (ThermoFisher) for RNA extraction. The extracted RNA was separated in a 15% Urea-TBE gel, and RNA fragments of 25 - 35 nt in length were excised for subsequent library construction.

Sequencing libraries were constructed following a previously established protocol (Ingolia et al., 2012) with modifications. Extracted RNA fragments were first treated with T4 Polynucleotide Kinase (T4 PNK, NEB, M0201L) without adding ATP to remove the 3’ end phosphates. A 5ʹ pre-adenylated and 3’ blocked adaptor (/5rApp/NNNNNNNNNNCACGGCGATCTTGCCGCC/3ddC/) was ligated to the 3’ ends of RNA fragments by using T4 RNA Ligase 2, truncated KQ (NEB, M0373L). The 5’ ends of RNA fragments were phosphorylated by T4 PNK and ligated to 5’ adaptor (CGATCTCCAATTCCCACTCCTTTCAAGACCTrC) using T4 RNA Ligase 1 (NEB, M0437M). Ribosomal RNA (rRNA) was depleted using RiboPOOL probes against human/mouse/rat rRNAs (siTOOLs BIOTECH). Reverse transcription (RT) was conducted using SuperScript IV Reverse Transcriptase (Thermo Fisher, 18090050) with RT primer GGGCGGCAAGATCGCCGTG. Libraries were amplified by PCR using Q5 Hot Start High-Fidelity DNA Polymerase (NEB, M0493L) with P5 primer (AATGATACGGCGACCACCGAG ATCTACACTCTTTCCCTACACGACGCTCTTCCGATCTNNAGGGCGGCAAGATCGCCG TG) and P7 primers (DMSO sample: CAAGCAGAAGACGGCATACGAGATCGGTTCAAGTGACTGGAGTTCAGACGTGTGCTCTTCCGATCTCCAATTCCCACTCCTTTCAAGACCT; Bicuculline sample: CAAGCAGAAGACGGCATACGAGATGCTGGATTGTGACTGGAGTTCAGACGTGTGCTCTTCCGATCTCCAATTCCCACTCCTTTCAAGACCT). After quality control, libraries were pooled and sequenced on a NovaSeq 6000 (Illumina) at the Yale Center for Genome Analysis.

### Ribosome profiling data analysis

After clipping both 5’ and 3’ adaptors, reads were deduplicated using FASTX-Toolkit. The 10 nt UMIs were then removed using Cutadapt. Reads were first mapped to a mammalian rRNA and tRNA reference using Bowtie. The remaining unmapped reads were then mapped to mouse reference genome GRCm39 and transcriptome by STAR. 28-31 nt footprints were used for subsequent analysis, with the P site offset for each footprint length being determined by using RiboMiner. For each read, P site was inferred by adding the P site offset to the 5’ coordinate.

## ACKNOWLEDGEMENT

We thank Susumu Tomita, Stephen Strittmatter, Arthur Horwich, Joan Steitz, Sreeganga Chandra, Thomas Biederer, Malaiyalam Mariappan, Yann Mineur, and members of the Guo lab for suggestions and comments on the manuscript, Carter Namkung and Ryan Stanton for assistance, and the Rodent Behavior Analysis facility supported by the Kavli Institute of Neuroscience, Yale Center for Genome Analysis, and Yale Genome Editing Center for technical support. This work was supported by an NIH Director’s New Innovator Award (DP2 GM132930) and by the National Institute of General Medical Sciences (R35 GM152208). P.L. and A.R.S. were supported by T32 NS041228 from the National Institute of Neurological Disorders and Stroke. M.R.P. and A.R.S. were supported by R01 MH077681 from the National Institute of Mental Health. J.U.G. is a New York Stem Cell Foundation−Robertson Investigator.

## SUPPLEMENTARY INFORMATION

**Supplementary Table 1** Predicted proteoform conversions by alternative translation initiation.

---

**Figure S1.**
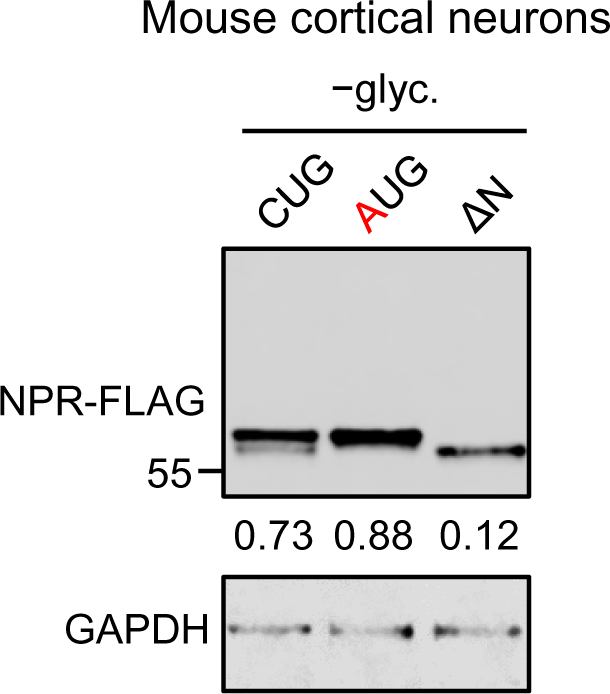
Alternative translation of NPR mRNA in primary neurons, related to Figure 1. Western blot analysis of wild-type (CUG) and mutant NPR cDNA in which CUG TIS was replaced with AUG TIS (AUG) or the first 11 codons were deleted (ΔN). Neurons were transduced with AAV for overexpression of NPR cDNA. Neuronal lysates were deglycosylated. Numbers indicate the fraction of long NPR.

**Figure S2.**
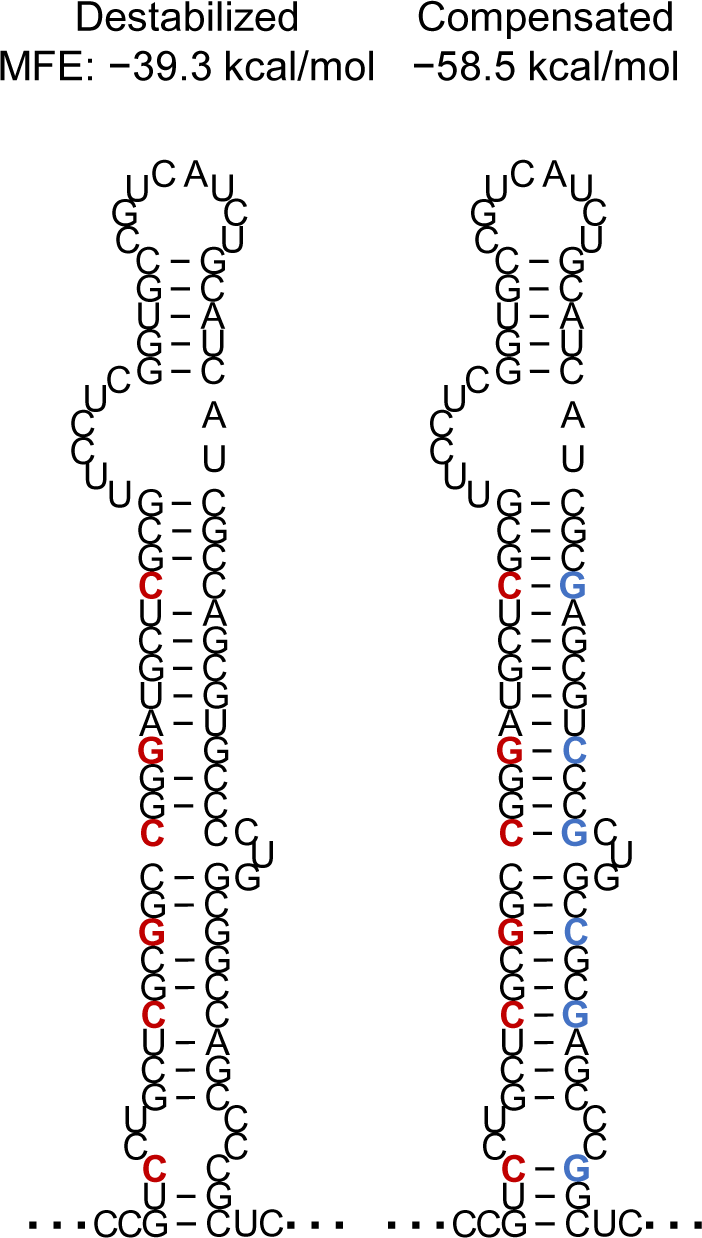
Predicted secondary structures of NPR mRNA mutants, related to Figure 2. The predicted secondary structure downstream of CUG TIS in NPR mRNA with mutations that disrupt or restore base-pairing. Destabilizing and compensatory mutations are indicated in red and blue, respectively. Predicted MFEs are shown.

**Figure S3.**
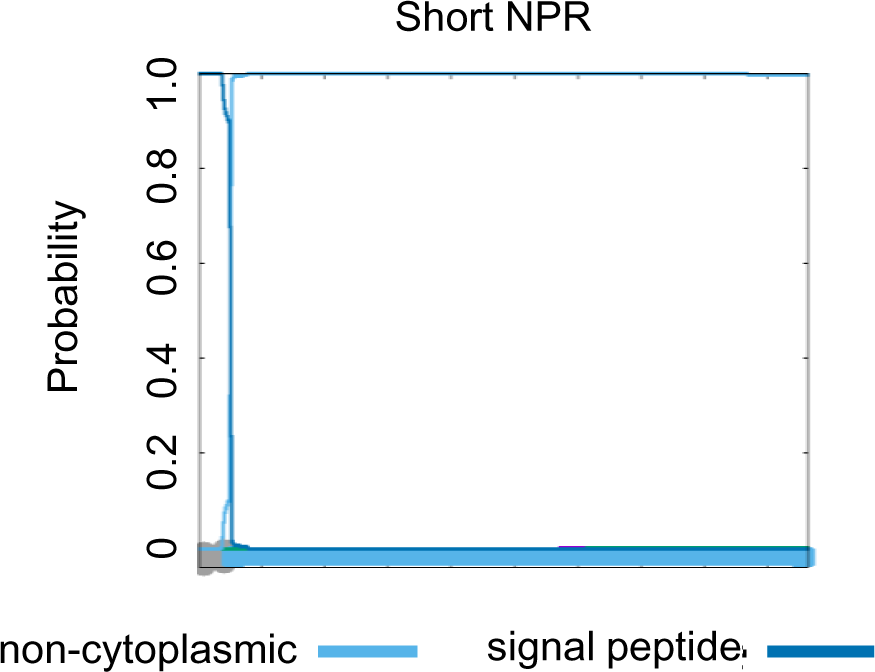
Phobius prediction of truncated N-terminal signal sequence acting as a cleavable signal peptide, related to Figure 3.

**Figure S4.**
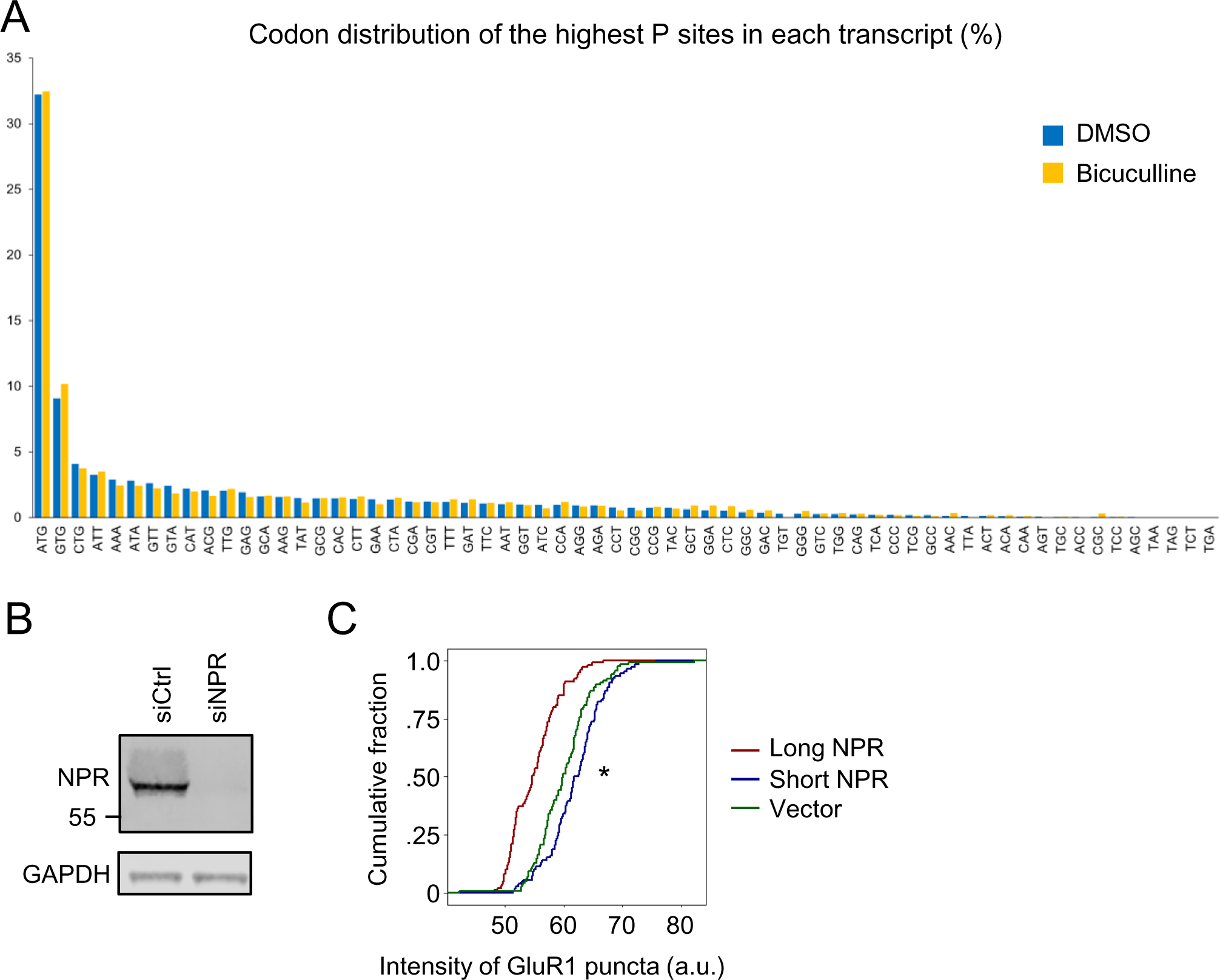
Distinct effects of long and short NPRs on synaptic AMPA receptor abundance, related to Figure 4. (**A**) Distribution of codons at the highest inferred P sites of initiating ribosome footprint in each transcript, sorted by percentage of each codon. (**B**) Western blot analysis of endogenous NPR in whole-cell lysates of cultured cortical neurons treated with either non-targeting siRNA (siCtrl) or siRNA against NPR (siNPR). (**C**) CDFs of surface GluR1 puncta mean intensity, comparing between neurons expressing either FLAG (vector), long, or short NPR in three independent experiments. \**P*=0.02 comparing the short NPR and vector groups by Kruskal−Wallis test.

**Figure S5.**
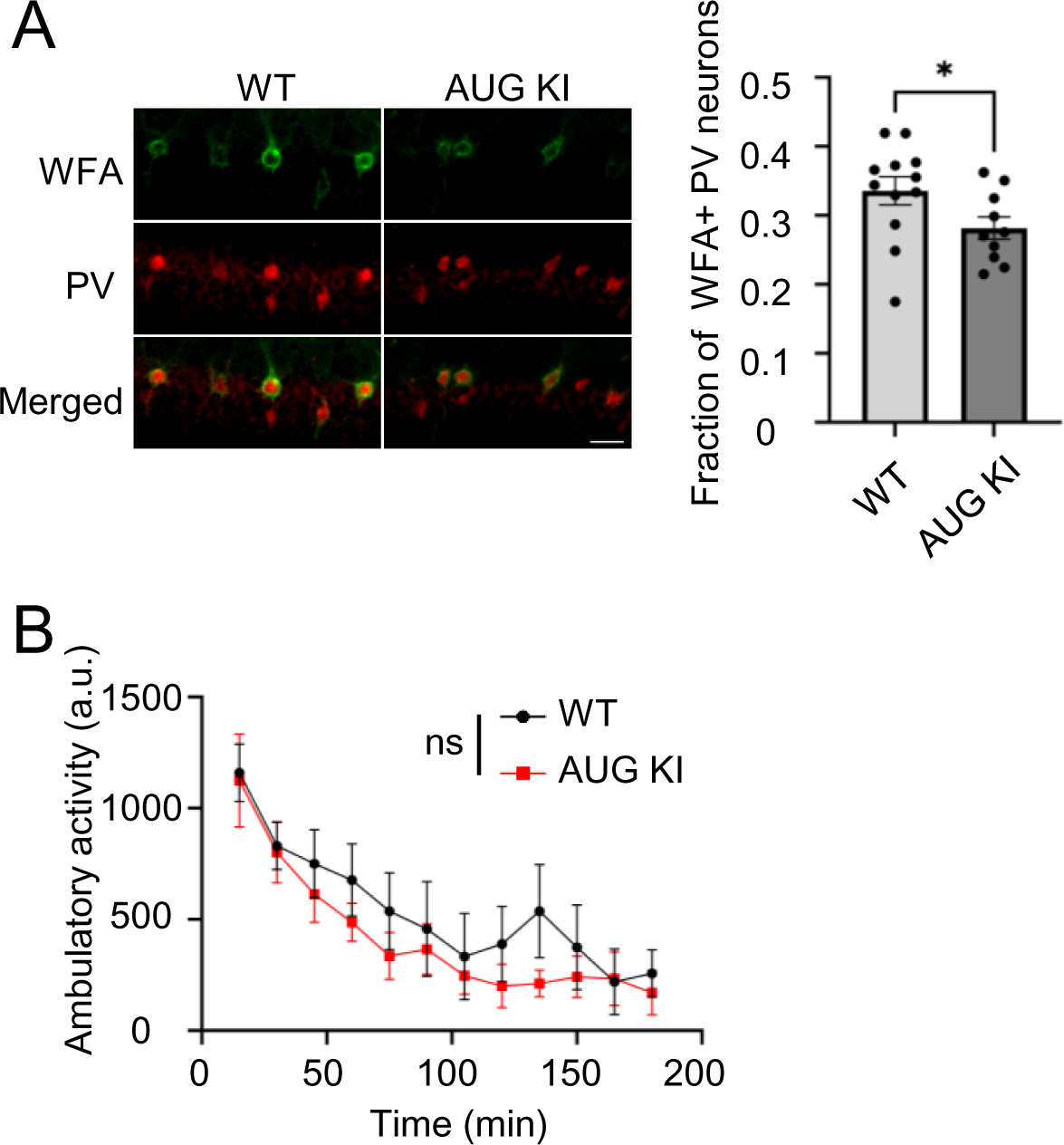
Additional characterization of NPTXR AUG KI mice, related to Figure 5. (**A**) Representative images (left) and quantification (right) of WFA and PV immunohistochemistry in CA1 regions of P30 WT and AUG KI mice (scale bar, 20 µm). *, *P*<0.05, Mann-Whitney test. (**B**) Spontaneous locomotor activity in a novel cage. n=5-6 animals per genotype. ns, *P>*0.05, 2-way Repeated Measures ANOVA.

**Figure S6.**
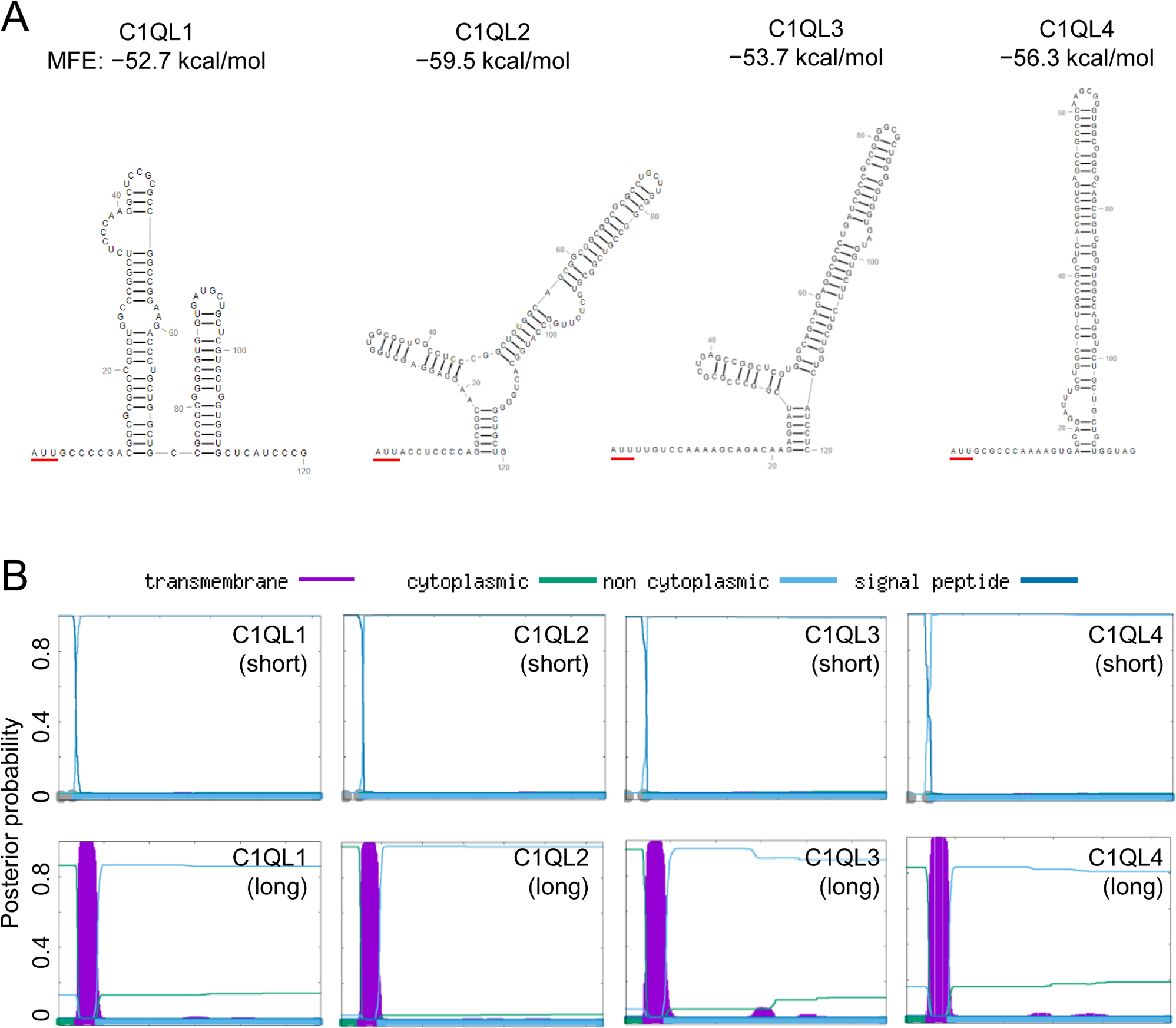
mRNA secondary structure and signal sequence predictions of C1QL1-4, related to Figure 6. (**A**) Predicted MFEs (top) of RNA secondary structures (bottom) downstream of AUU TIS in C1QL1-4 mRNAs. (**B**) Phobius predictions of signal peptide sequences and transmembrane domains of short and long proteoforms of C1QL1-4.

## REFERENCE

Atkins, J.F., Loughran, G., Bhatt, P.R., Firth, A.E., and Baranov, P.V. (2016). Ribosomal frameshifting and transcriptional slippage: From genetic steganography and cryptography to adventitious use. Nucleic Acids Res 44, 7007–7078. 10.1093/nar/gkw530.

Bogaert, A., Fernandez, E., and Gevaert, K. (2020). N-Terminal Proteoforms in Human Disease. Trends Biochem Sci 45, 308–320. 10.1016/j.tibs.2019.12.009.

Bolliger, M.F., Martinelli, D.C., and Sudhof, T.C. (2011). The cell-adhesion G protein-coupled receptor BAI3 is a high-affinity receptor for C1q-like proteins. Proc Natl Acad Sci U S A 108, 2534–2539. 10.1073/pnas.1019577108.

Calligaris, R., Bottardi, S., Cogoi, S., Apezteguia, I., and Santoro, C. (1995). Alternative translation initiation site usage results in two functionally distinct forms of the GATA-1 transcription factor. Proc Natl Acad Sci U S A 92, 11598–11602. 10.1073/pnas.92.25.11598.

Chang, M.C., Park, J.M., Pelkey, K.A., Grabenstatter, H.L., Xu, D., Linden, D.J., Sutula, T.P., McBain, C.J., and Worley, P.F. (2010). Narp regulates homeostatic scaling of excitatory synapses on parvalbumin-expressing interneurons. Nat Neurosci 13, 1090–1097. 10.1038/nn.2621.

Chen, J., Brunner, A.D., Cogan, J.Z., Nunez, J.K., Fields, A.P., Adamson, B., Itzhak, D.N., Li, J.Y., Mann, M., Leonetti, M.D., and Weissman, J.S. (2020). Pervasive functional translation of noncanonical human open reading frames. Science 367, 1140–1146. 10.1126/science.aay0262.

Cho, R.W., Park, J.M., Wolff, S.B., Xu, D., Hopf, C., Kim, J.A., Reddy, R.C., Petralia, R.S., Perin, M.S., Linden, D.J., and Worley, P.F. (2008). mGluR1/5-dependent long-term depression requires the regulated ectodomain cleavage of neuronal pentraxin NPR by TACE. Neuron 57, 858–871. 10.1016/j.neuron.2008.01.010.

Dodds, D.C., Omeis, I.A., Cushman, S.J., Helms, J.A., and Perin, M.S. (1997). Neuronal pentraxin receptor, a novel putative integral membrane pentraxin that interacts with neuronal pentraxin 1 and 2 and taipoxin-associated calcium-binding protein 49. J Biol Chem 272, 21488–21494. 10.1074/jbc.272.34.21488.

Gao, X., Wan, J., Liu, B., Ma, M., Shen, B., and Qian, S.B. (2015). Quantitative profiling of initiating ribosomes in vivo. Nat Methods 12, 147–153. 10.1038/nmeth.3208.

Gruber, A.R., Lorenz, R., Bernhart, S.H., Neubock, R., and Hofacker, I.L. (2008). The Vienna RNA websuite. Nucleic Acids Res 36, W70–74. 10.1093/nar/gkn188.

Ingolia, N.T., Brar, G.A., Rouskin, S., McGeachy, A.M., and Weissman, J.S. (2012). The ribosome profiling strategy for monitoring translation in vivo by deep sequencing of ribosome-protected mRNA fragments. Nat Protoc 7, 1534–1550. 10.1038/nprot.2012.086.

Ingolia, N.T., Lareau, L.F., and Weissman, J.S. (2011). Ribosome profiling of mouse embryonic stem cells reveals the complexity and dynamics of mammalian proteomes. Cell 147, 789–802. 10.1016/j.cell.2011.10.002.

Ivanov, I.P., Firth, A.E., Michel, A.M., Atkins, J.F., and Baranov, P.V. (2011). Identification of evolutionarily conserved non-AUG-initiated N-terminal extensions in human coding sequences. Nucleic Acids Res 39, 4220–4234. 10.1093/nar/gkr007.

Kall, L., Krogh, A., and Sonnhammer, E.L. (2004). A combined transmembrane topology and signal peptide prediction method. J Mol Biol 338, 1027–1036. 10.1016/j.jmb.2004.03.016.

Kall, L., Krogh, A., and Sonnhammer, E.L. (2007). Advantages of combined transmembrane topology and signal peptide prediction--the Phobius web server. Nucleic Acids Res 35, W429–432. 10.1093/nar/gkm256.

Kapp, L.D., and Lorsch, J.R. (2004). The molecular mechanics of eukaryotic translation. Annu Rev Biochem 73, 657–704. 10.1146/annurev.biochem.73.030403.080419.

Kazak, L., Reyes, A., Duncan, A.L., Rorbach, J., Wood, S.R., Brea-Calvo, G., Gammage, P.A., Robinson, A.J., Minczuk, M., and Holt, I.J. (2013). Alternative translation initiation augments the human mitochondrial proteome. Nucleic Acids Res 41, 2354–2369. 10.1093/nar/gks1347.

Kozak, M. (1990). Downstream secondary structure facilitates recognition of initiator codons by eukaryotic ribosomes. Proc Natl Acad Sci U S A 87, 8301–8305. 10.1073/pnas.87.21.8301.

Lak, B., Li, S., Belevich, I., Sree, S., Butkovic, R., Ikonen, E., and Jokitalo, E. (2021). Specific subdomain localization of ER resident proteins and membrane contact sites resolved by electron microscopy. Eur J Cell Biol 100, 151180. 10.1016/j.ejcb.2021.151180.

Lee, S., Liu, B., Lee, S., Huang, S.X., Shen, B., and Qian, S.B. (2012). Global mapping of translation initiation sites in mammalian cells at single-nucleotide resolution. Proc Natl Acad Sci U S A 109, E2424–2432. 10.1073/pnas.1207846109.

Lee, S.J., Wei, M., Zhang, C., Maxeiner, S., Pak, C., Calado Botelho, S., Trotter, J., Sterky, F.H., and Sudhof, T.C. (2017). Presynaptic Neuronal Pentraxin Receptor Organizes Excitatory and Inhibitory Synapses. J Neurosci 37, 1062–1080. 10.1523/JNEUROSCI.2768-16.2016.

Lin, M.F., Jungreis, I., and Kellis, M. (2011). PhyloCSF: a comparative genomics method to distinguish protein coding and non-coding regions. Bioinformatics 27, i275–282. 10.1093/bioinformatics/btr209.

Loughran, G., Chou, M.Y., Ivanov, I.P., Jungreis, I., Kellis, M., Kiran, A.M., Baranov, P.V., and Atkins, J.F. (2014). Evidence of efficient stop codon readthrough in four mammalian genes. Nucleic Acids Res 42, 8928–8938. 10.1093/nar/gku608.

Martinelli, D.C., Chew, K.S., Rohlmann, A., Lum, M.Y., Ressl, S., Hattar, S., Brunger, A.T., Missler, M., and Sudhof, T.C. (2016). Expression of C1ql3 in Discrete Neuronal Populations Controls Efferent Synapse Numbers and Diverse Behaviors. Neuron 91, 1034–1051. 10.1016/j.neuron.2016.07.002.

Matsuda, K., Budisantoso, T., Mitakidis, N., Sugaya, Y., Miura, E., Kakegawa, W., Yamasaki, M., Konno, K., Uchigashima, M., Abe, M., et al. (2016). Transsynaptic Modulation of Kainate Receptor Functions by C1q-like Proteins. Neuron 90, 752–767. 10.1016/j.neuron.2016.04.001.

Murray, A.J., Woloszynowska-Fraser, M.U., Ansel-Bollepalli, L., Cole, K.L., Foggetti, A., Crouch, B., Riedel, G., and Wulff, P. (2015). Parvalbumin-positive interneurons of the prefrontal cortex support working memory and cognitive flexibility. Sci Rep 5, 16778. 10.1038/srep16778.

Park, J., Chavez, A.E., Mineur, Y.S., Morimoto-Tomita, M., Lutzu, S., Kim, K.S., Picciotto, M.R., Castillo, P.E., and Tomita, S. (2016). CaMKII Phosphorylation of TARPgamma-8 Is a Mediator of LTP and Learning and Memory. Neuron 92, 75–83. 10.1016/j.neuron.2016.09.002.

Pelkey, K.A., Barksdale, E., Craig, M.T., Yuan, X., Sukumaran, M., Vargish, G.A., Mitchell, R.M., Wyeth, M.S., Petralia, R.S., Chittajallu, R., et al. (2015). Pentraxins coordinate excitatory synapse maturation and circuit integration of parvalbumin interneurons. Neuron 85, 1257–1272. 10.1016/j.neuron.2015.02.020.

Schaukowitch, K., Reese, A.L., Kim, S.K., Kilaru, G., Joo, J.Y., Kavalali, E.T., and Kim, T.K. (2017). An Intrinsic Transcriptional Program Underlying Synaptic Scaling during Activity Suppression. Cell Rep 18, 1512–1526. 10.1016/j.celrep.2017.01.033.

Shatsky, I.N., Terenin, I.M., Smirnova, V.V., and Andreev, D.E. (2018). Cap-Independent Translation: What’s in a Name? Trends Biochem Sci 43, 882–895. 10.1016/j.tibs.2018.04.011.

Tsang, M.J., and Cheeseman, I.M. (2023). Alternative CDC20 translational isoforms tune mitotic arrest duration. Nature 617, 154–161. 10.1038/s41586-023-05943-7.

Tsui, C.C., Copeland, N.G., Gilbert, D.J., Jenkins, N.A., Barnes, C., and Worley, P.F. (1996). Narp, a novel member of the pentraxin family, promotes neurite outgrowth and is dynamically regulated by neuronal activity. J Neurosci 16, 2463–2478. 10.1523/JNEUROSCI.16-08-02463.1996.

Van Damme, P., Gawron, D., Van Criekinge, W., and Menschaert, G. (2014). N-terminal proteomics and ribosome profiling provide a comprehensive view of the alternative translation initiation landscape in mice and men. Mol Cell Proteomics 13, 1245–1261. 10.1074/mcp.M113.036442.

Wang, J., Shin, B.S., Alvarado, C., Kim, J.R., Bohlen, J., Dever, T.E., and Puglisi, J.D. (2022). Rapid 40S scanning and its regulation by mRNA structure during eukaryotic translation initiation. Cell 185, 4474–4487 e4417. 10.1016/j.cell.2022.10.005.

Williams, C.C., Jan, C.H., and Weissman, J.S. (2014). Targeting and plasticity of mitochondrial proteins revealed by proximity-specific ribosome profiling. Science 346, 748–751. 10.1126/science.1257522.

Wu, B., Eliscovich, C., Yoon, Y.J., and Singer, R.H. (2016). Translation dynamics of single mRNAs in live cells and neurons. Science 352, 1430–1435. 10.1126/science.aaf1084.

Xu, D., Hopf, C., Reddy, R., Cho, R.W., Guo, L., Lanahan, A., Petralia, R.S., Wenthold, R.J., O’Brien, R.J., and Worley, P. (2003). Narp and NP1 form heterocomplexes that function in developmental and activity-dependent synaptic plasticity. Neuron 39, 513–528. 10.1016/s0896-6273(03)00463-x.

Yau, J.O., Chaichim, C., Power, J.M., and McNally, G.P. (2021). The Roles of Basolateral Amygdala Parvalbumin Neurons in Fear Learning. J Neurosci 41, 9223–9234. 10.1523/JNEUROSCI.2461-20.2021.

